# Autonomous oscillations and phase-locking in a biophysically detailed model of the STN-GPe network

**DOI:** 10.1101/611103

**Authors:** Lucas A. Koelman, Madeleine M. Lowery

## Abstract

The aim of this study was to understand the relative role of autonomous oscillations and patterning by exogenous oscillatory inputs in the generation of pathological oscillatory activity within the subthalamic nucleus (STN) - external globus pallidus (GPe) network in Parkinson’s disease. A biophysically detailed model that accounts for the integration of synaptic currents and their interaction with intrinsic membrane currents in dendritic structures within the STN and GPe was developed. The model was used to investigate the development of beta-band synchrony and bursting within the STN-GPe network by changing the balance of excitation and inhibition in both nuclei, and by adding exogenous oscillatory inputs with varying phase relationships through the hyperdirect cortico-subthalamic and indirect striato-pallidal pathways. The model showed an intrinsic susceptibility to beta-band oscillations that was manifest in weak autonomously generated oscillations within the STN-GPe network and in selective amplification of exogenous beta-band synaptic inputs near the network’s endogenous oscillation frequency. The resonant oscillation frequency was determined by the net level of excitatory drive in the loop. Intrinsically generated oscillations were too weak to support a pacemaker role for the STN-GPe network, however, they were considerably amplified by sparse cortical beta inputs when their frequency range overlapped and were further amplified by striatal beta inputs that promoted anti-phase firing of the cortex and GPe, resulting in maximum transient inhibition of STN neurons. The model elucidates a mechanism of cortical patterning of the STN-GPe network through feedback inhibition whereby intrinsic susceptibility to beta-band oscillations can lead to phase locked spiking under parkinsonian conditions. These results point to resonance of endogenous oscillations with exogenous patterning of the STN-GPe network as a mechanism of pathological synchronization, and a role for the pallido-striatal feedback loop in amplifying beta oscillations.

**Author summary:** Exaggerated beta-frequency neuronal synchrony is observed throughout the basal ganglia in Parkinson’s disease and is reduced with medication and during deep brain stimulation. The power of beta-band oscillations is increasingly used as a biomarker to guide antiparkinsonian therapies. Despite their importance as a clinical target, the mechanisms by which pathological beta-band oscillations are generated are not yet clearly understood. In vitro electrophysiological recordings support a theory of enhanced phase locking of the reciprocally connected subthalamo-pallidal network to beta-band cortical inputs but this has not yet been clearly demonstrated in a model. We present a new model of the subthalamo-pallidal network consisting of biophysically detailed cell models that captures the interaction between synaptic and intrinsic currents in dendritic structures. The model shows how phase locking of subthalamic and pallidal neurons and exaggerated bursting in subthalamic neurons can arise from the interaction of these currents when the balance of excitation and inhibition is changed and how phase locking is amplified under specific phase relationships between cortical and striatal beta inputs.

## Introduction

Pathological oscillations in the basal ganglia-thalamocortical (BGTC) network have long been implicated in the motor symptoms of Parkinson’s disease. Beta-band (15-30 Hz) oscillations are strengthened consistently after dopamine depletion (DD) both in Parkinson’s disease (PD) patients and parkinsonian animal models [1–3], and are reduced by deep brain stimulation and pharmacological interventions that alleviate parkinsonian motor symptoms [4–7]. The magnitude of beta oscillations has also been shown to correlate with the severity and degree of improvement of bradykinetic/akinetic motor symptoms and rigidity [4, 8].

The reciprocally connected subthalamo-pallidal network is a key site in the basal ganglia (BG) in which beta-band oscillations are manifest in Parkinson’s disease [2, 9]. It is not clear whether it plays an active part in generating beta-band oscillations, nor whether it amplifies or merely sustains them. Neither is it fully understood how beta-band oscillations relate to other pathological patterns of neural activity in the subthalamic nucleus (STN) and external globus pallidus (GPe) that correlate more strongly with parkinsonian motor symptoms, notably increased neural bursting [10, 11]. It is clear, however, that interventions in the loop and its afferents that reduce beta-band oscillations [12] or bursting [13–15], in the form of precise chemical lesions or optogenetic stimulation targeted at the subcellular level, lead to improvements in motor symptoms. Similarly, the STN and GPe are effective targets for deep brain stimulation (DBS) [16, 17], which likely activates a more diverse set of neuronal elements [18]. Although at present it is unknown whether a causal relationship exists between beta-frequency activity and parkinsonian motor symptoms, due to the strong correlation between the two it is important to understand the biophysical mechanisms underlying these pathological activity patterns.

Computational models provide a valuable tool with which to explore different hypotheses regarding the origin of pathological synchronization in the BG and their suppression by means of therapeutic interventions. The most prominent hypothesis generally emphasize the importance of a particular feedback loop between subpopulations of cells within the multiple interconnected loops that constitute the full BGTC network. Different models have placed the origin of beta and sub-beta band oscillations in the STN-GPe network [19–22], in striatal or pallidostriatal circuits [23, 24], or in the full BGTC loop [25–28] (reviewed in [29, 30]). These models, based on experimentally observed oscillations in different subcircuits of the BG, show that many of them are prone to oscillate, be it through intrinsic pacemaking or susceptibility to an extrinsic rhythm.

The STN-GPe network was an early focus of modelling studies due to its reciprocally connected structure and because it can generate low frequency oscillations autonomously when isolated in tissue cultures [31]. Models of the STN-GPe as a pacemaker initially focused on the generation of low frequency oscillations within the frequency range of parkinsonian tremor [19, 32], with focus shifting to the beta-band with increasing evidence of a correlation between beta activity and bradykinesia and akinesia [21, 22]. More recent experimental evidence suggests that, rather than it operating in a pacemaking mode, patterning by cortex may play a critical role in the generation of beta-band oscillations in the STN-GPe network in Parkinson’s disease. This is supported by observations of high functional coupling between cortex and STN [1, 9, 33–36], and that oscillatory activity in STN-GPe is contingent on input from cortex and can be abolished by disrupting these inputs [12, 37, 38].

Cortical patterning of the STN-GPe network by means of feedback inhibition provides a proposed mechanism for this functional coupling [9, 12, 39–41]. In this hypothesis, weak oscillatory activity arriving via cortico-STN afferents is amplified in the STN-GPe network when feedback inhibition by GPe is offset in phase to cortical excitation. While such feedback-mediated oscillations have been observed in vivo [39, 42], the ability of the network to generate autonomous oscillations and its resonant response properties are still poorly understood.

To explain oscillatory properties, previous modeling studies have focused on alterations in connection patterns and strengths within or between nuclei, typically represented by mean-field or single-compartment spiking neuron models. While such models are computationally efficient, they may not fully capture the role of intrinsic properties of neurons in shaping pathological activity patterns (reviewed in [30]). Although cell-specific ion channels can be used, single-compartment neuron models lump together ion channels and synapses in one isopotential compartment in a way that may not capture the complex dynamics that arise when non-uniformly distributed ion channels [43] interact with synapses associated with distinct subcellular regions [15, 44, 45]. Hence they may not fully account for the mechanisms contributing to cortical patterning and the role that synaptic-ionic current interactions play in sustaining beta-band oscillations and excessive burst firing.

Recent evidence suggests that following dopamine depletion the balance of excitatory and inhibitory synaptic currents in STN neurons is shifted toward inhibition [46, 47] which is known to promote burst responses through increasing the availability of *Ca*^2+^ and *Na*^+^ channels that are de-inactivated at hyperpolarized potentials [39]. In the GPe an increase in inhibition, caused mainly by strengthening of striato-pallidal afferents, is also believed to play a role in generating pathological oscillations as demonstrated in model simulation [19, 21, 32, 48]. Based on experimental data, increased GPe inhibition has been suggested to cause increased engagement of HCN channels [49], which are involved in phase resetting and controlling the regularity of firing. However, whether the functional coupling between BG nuclei is also moderated by the excitation-inhibition balance is not fully understood.

The aim of this study was, therefore, to understand the conditions under which cortical patterning of the STN-GPe network arises in a biophysically plausible manner and how this is related to the balance of excitation and inhibition in each population. The STN-GPe network was modeled using biophysically detailed multi-compartmental cell models of STN and GPe neurons that capture the interaction between synaptic and intrinsic currents distributed within the dendritic structure and involved in autonomous pacemaking and bursting [43, 50] (Fig. 1). The generation of oscillations both autonomously within the network and in response to beta frequency inputs from the cortex (CTX) and striatal medium spiny neurons (MSN) was examined as the balance of excitation and inhibition within the network was systematically varied, and oscillatory inputs with varying phase relationships were added. A better understanding of the relative contribution of these different factors and their interaction has the potential to improve understanding of the mechanism of action of existing anti-parkinsonian therapies, including deep brain stimulation (DBS) and to guide the development of more effective circuit interventions.

**Fig 1.**
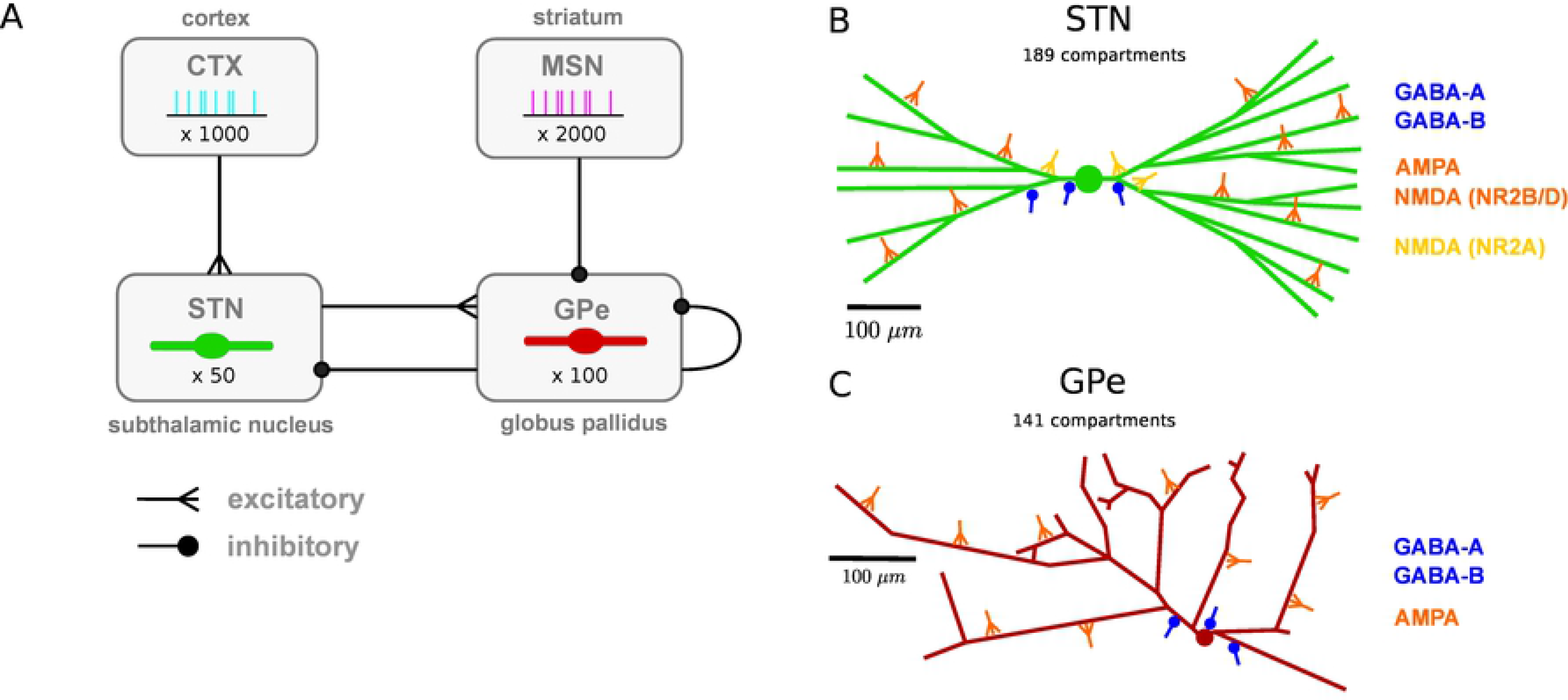
Network architecture: population and subcellular connectivity. **A**: Neuronal populations and their projections modeled in the network. Subthalamic projection neurons (STN) and prototypic neurons of the external globus pallidus (GPe) were modeled using multi-compartmental neuron models. Cortical projection neurons (CTX) and striatal medium spiny neurons (MSN) were modeled as spike generators. **B**: Branching structure of the STN neuron model and representative synapses by afferent type, indicating subcellular distribution of synapses. Cortical glutamergic afferents synapse primarily onto thin dendrites, distally relative to the soma, but NMDA receptors with faster NR2A subunits mainly target the soma and proximal areas. Pallidal GABAergic afferents target proximal areas of the cell. **C**: Branching structure of the GPe neuron model and representative synapses by afferent type. GABAergic GPe-GPe collaterals mainly target somata and proximal dendrites, whereas glutamergic afferents were placed in distal regions. Full details of the model are provided in the Methods.

## Results

The STN-GPe pacemaker hypothesis was first investigated by modeling cortical inputs to the STN as Poisson spike generators without any periodic or oscillatory component (Sec. 4.1-4.3). Cortical patterning of neural activity in the STN-GPe network via the hyperdirect pathway was then investigated by modeling cortical input to the STN inputs as periodically bursting spike trains. To investigate whether changing the excitation-inhibition balance in STN and GPe contributed to changes in spontaneous synchronization and functional coupling between nuclei, we shifted the ratio of excitation and inhibition by systematically altering the strength of individual projections between nuclei. The ratio of total excitatory to inhibitory synaptic currents (E/I ratio) was shifted by scaling the peak conductance of all synapses belonging to a given projection known to be strengthened or weakened by dopamine depletion. The role of additional oscillatory inputs entering the STN-GPe network via the indirect striato-pallidal pathway and their phase relationship to cortical inputs was then investigated.

### The excitation-inhibition balance in the STN sets the oscillation frequency of the STN-GPe network and firing mode of STN neurons

Sweeping the synaptic conductance of cortico-STN afferents moved the STN-GPe network through a diverse range of firing patterns (Fig 2). Increasing the synaptic conductance of cortical Poisson-distributed inputs caused a proportional increase in net excitatory current to the STN resulting in increased excitatory input from STN to GPe. However, the GPe population exhibited a saturating population firing rate curve due to two negative feedback mechanisms that counteract the effect of increasing STN excitation and have a homeostatic effect on the GPE’s ratio of excitation to inhibition: short-term depression of STN to GPe synapses [51] and reciprocal inhibition through intra-GPe colaterals. As a result, the increase in feedback inhibition from the GPe did not keep pace with the rise in excitation from the cortex (Fig 2.Ai-ii). At the lower end of the synaptic conductance scale, both excitation and inhibition were low and the ratio of excitatory to inhibitory synaptic currents (E/I ratio) was balanced towards inhibition (E/I < 1). STN and GPe neurons showed high spike train synchronization at 12 Hz (population vector length of 0.53; 0.68 in STN; GPe, conditioned on GPe phase) with spike trains consisting largely of singlets and doublets. The STN population vector led that of GPe by approximately 45 degrees indicating that STN neurons spike at the end of an inhibitory episode and then trigger a wave of GPe spikes approximately 10 ms later.

**Fig 2.**
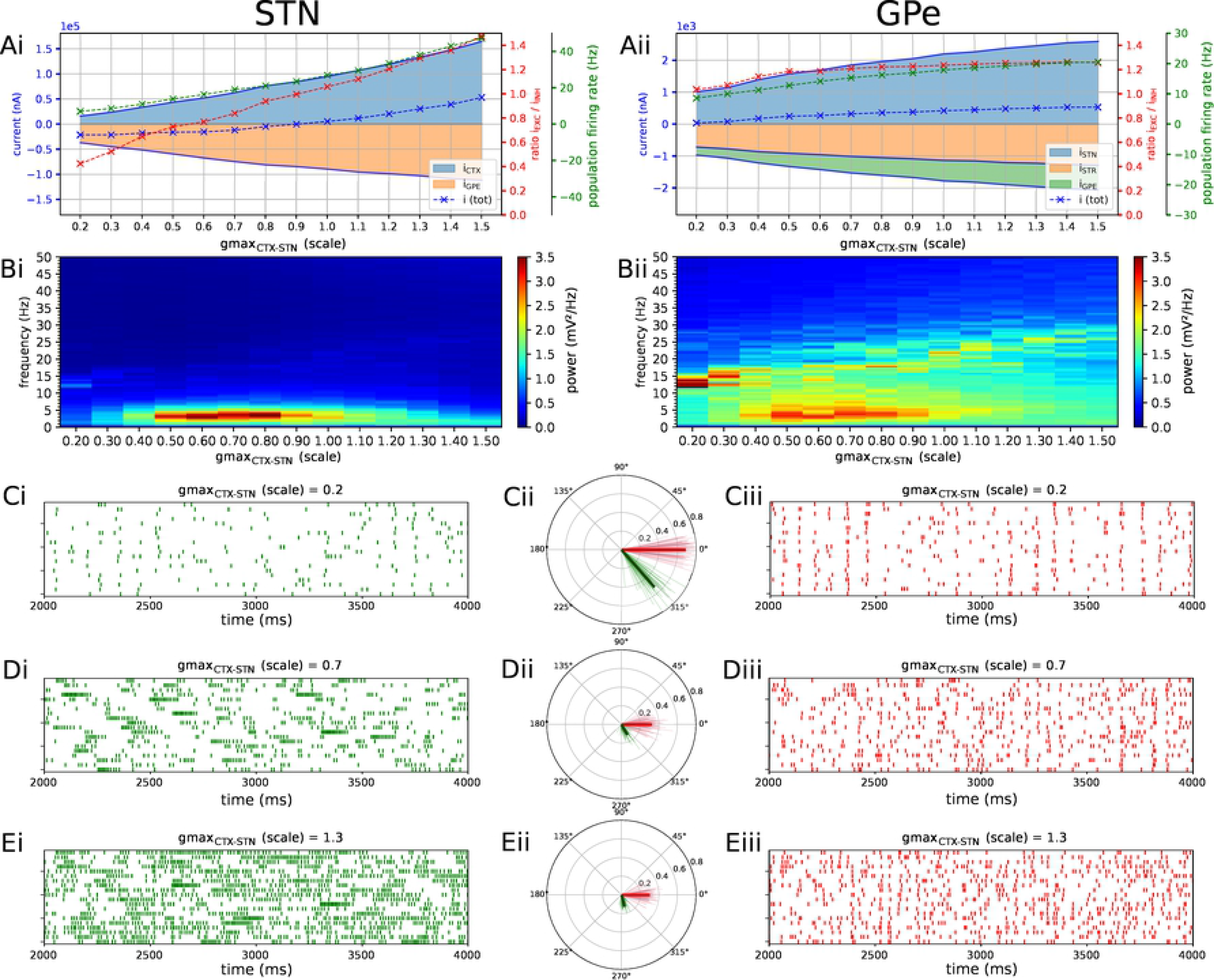
The level of excitation by cortex determines firing patterns and oscillation frequency in the autonomous STN-GPe network. Characterization of activity in the autonomous STN-GPe network for increasing values of the CTX to STN synaptic conductance. **A**: Population firing rate (green), E/I ratio (red), and net synaptic current (blue) in STN (Ai) and GPe (Aii). Shaded areas represent estimated total synaptic current from one pre-synaptic population during a simulation. Total current was estimated by recording all synapses on 3 randomly selected cells in each population and adjusting for the true number of cells. **B**: Mean PSD of the somatic membrane voltages of STN (Bi) and GPe (Bii) neurons, showing the slow bursting regimen for intermediate excitation-inhibition balance in STN. Beta-band oscillation was not reliably transmitted to STN when slow GABA_B_R-mediated synaptic currents dominated over fast-acting GABA_A_ currents. **C-E**: Representative spike trains and phase vectors for STN (column i, green) and GPe population (column iii, red) for three different scale factors of the CTX to STN conductance (scale 0.2; 0.7; 1.3 in rows C; D; E respectively). Column ii shows phase vectors of the STN and GPe populations (in green; red, respectively, mean population vectors plotted as thick solid lines and cell vectors as thin transparent lines) reflecting phase locking to the instantaneous GPe phase. Phase vectors were conditioned on the instantaneous phase of the GPe population extracted using a bandpass filter with passband of 8 Hz centered on the frequency bin with maximum power in the 13-30 Hz band.

The synchronization frequency of GPe neurons increased across the parameter sweep but oscillation power was low and the oscillation was weakly transmitted to STN neurons in the baseline model (Fig 2.Bi-Bii). Although synchronization of STN neurons was weak throughout most of the parameter range, spikes showed a tendency to occur in a fixed phase relationship with GPe as evidenced by the alignment of individual and population phase vectors (Fig 2.Cii,Dii,Eii) and by the phase histogram (Fig 5.Bi). Only when both the excitatory and inhibitory drive to STN were low, and there was a net inhibition to STN neurons, did the STN and GPe synchronize strongly (Fig 2.Bi-Bii, gmax scale = 0.2; Fig 2.Ci-Ciii). This was because in the baseline model, GPe to STN inhibition was dominated by slow GABA_B_ receptor (GABA_B_R) mediated synaptic currents that promote low-frequency bursting and counteract faster synchronization in STN. These slow GABA_B_R-mediated currents are facilitated by faster pre-synaptic spiking in a near-sigmoidal fashion and thus tend to dominate at higher GPe firing rates. When GPe to STN inhibition was shifted to the faster GABA_A_ synapses, low-frequency bursting was reduced, beta-band synchronization was strengthened and the frequency was strongly determined by the level of cortical excitation (Fig 5.E). In the intermediate range of the cortical synapse strength (0.5 ≤ scale ≤ 1.0), the total current from both excitatory CTX inhibitory and GPe afferents increased but excitation outpaced inhibition because of the negative feedback mechanisms in the GPe described above. Excitation and inhibition were relatively balanced (0.7 ≤ *E/I* ≤ 1.1) but the total inhibitory current was higher. As detailed above, higher inhibition in the dendrites leads to higher availability of dendritic ion channels underlying burst responses. Hence, the shift to higher inhibition brought STN neurons into a slow burst firing mode with sparse, strong bursts with higher intraburst firing rate (Fig 2.Dii). These low-frequency fluctuations in firing rate were transmitted to GPe neurons and showed up clearly in the power spectrum of both nuclei (Fig 2.Bi-Bii). As the CTX to STN conductance was increased further, resulting in a net excitation to STN, STN firing rates increased accordingly and became more uniform, reflected in a lower coefficient of variation and a convergence of the background and intra-burst firing rates (Fig 2.Ei).

Increasing the synaptic conductance of the inhibitory GPe-GPe projection moved the STN-GPe network through a similar trajectory of firing patterns while shifting the excitation-inhibition balance in STN and GPe in opposing directions (Fig. 3). The strong low-frequency burst firing mode of STN cells was engaged within the same interval of E/I ratios (Fig. 3, Ai, Bi, F), and firing became more temporally uniform as this ratio increased to favour excitation (Fig. 3, Ci, Di, Eii, F). As with the parameter sweep of the CTX-STN conductance, the mean firing rate of the GPe population was less sensitive to the synaptic conductance than that of the STN population. The negative feedback structure of the STN-GPe network was engaged to maintain the mean GPe firing rate within a narrow range: the increase in collateral GPe-GPe inhibition was offset by the increased excitation by STN cells as they were disinhibited (Fig. 3.Aii). GPe neurons again synchronized more strongly than STN neurons: the 13-30 Hz peak was prominent in its power spectral density (PSD) (Fig. 3.B) and single unit spikes were more tightly locked to the phase of the oscillation compared to STN (this was true whether the instantaneous phase was extracted from the STN or GPe population, data not shown). Although STN cells were overall more likely to spike during a specific phase of the oscillation, their tendency to emit longer bursts rather than short bursts or singlets/doublets reduced the precision of phase locking and, therefore, their phase vector length compared to GPe cells (Fig. 3.C-D). The oscillation frequency in GPe shifted downward as the GPe-GPe conductance was decreased and the excitatory drive from GPe decreased accordingly (Fig. 3.Bii), with a near-merging of the sub-beta and beta-range peaks in the lower conductance range.

**Fig 3.**
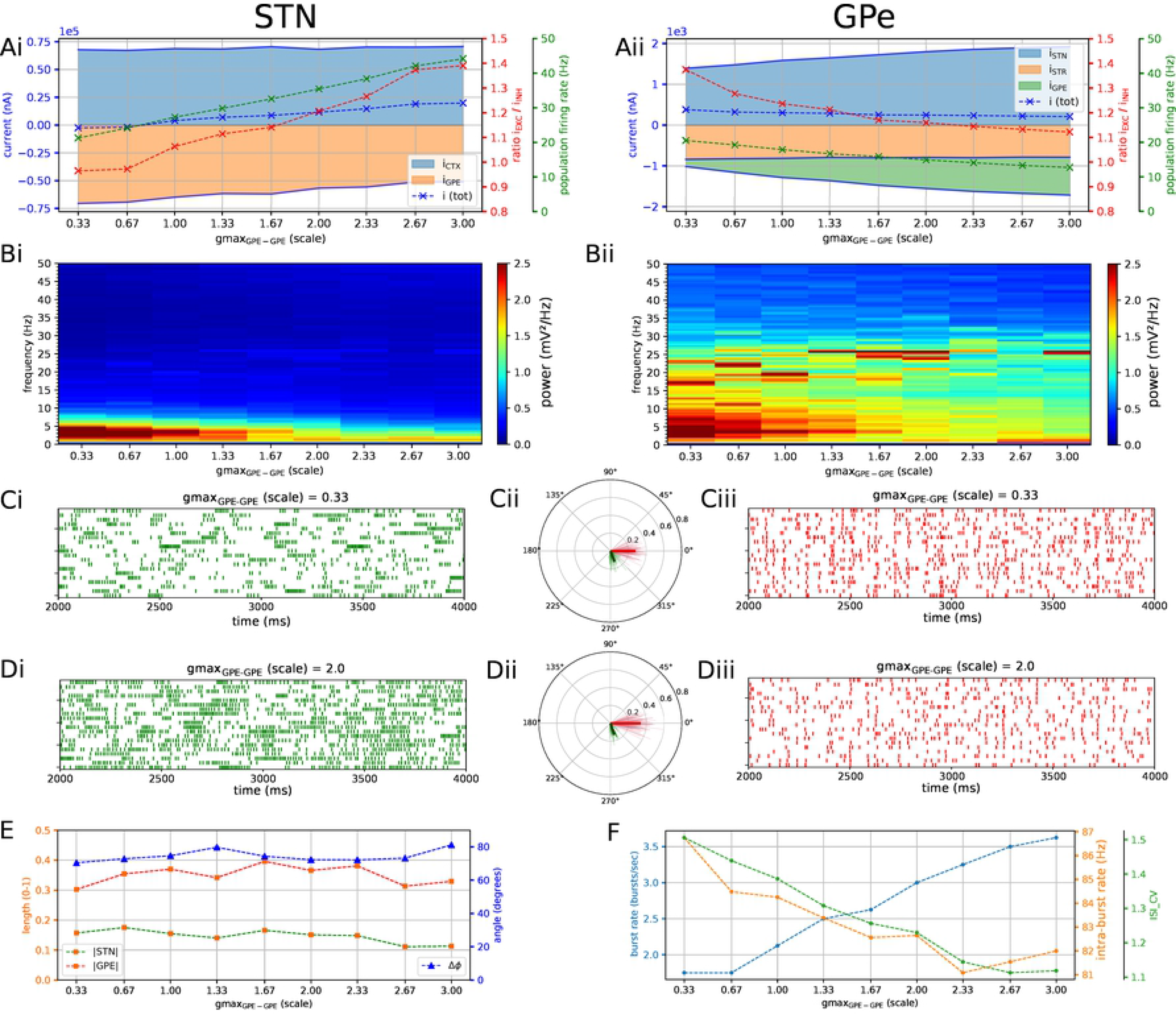
Increasing the level of collateral GPe-GPe inhibition shifts the excitation-inhibition balance in STN and GPe in opposite directions. Characterization of activity in the autonomous STN-GPe network for increasing values of GPe-GPe synaptic conductance. **A**: Population firing rate (green), E/I ratio (red), and net synaptic current (blue) in STN (Ai) and GPe (Aii). Shaded areas represent estimated total synaptic current from one pre-synaptic population during a simulation. **B**: Mean PSD of the somatic membrane voltages of STN (Bi) and GPe (Bii) neurons. Low-frequency bursting in STN neurons is engaged for weak GPe-GPe inhibition leading to strong GPe to STN inhibition (Bi). GPe neurons synchronized at beta frequencies (Bii). **C-D**: Representative spike trains and phase vectors for STN (column i, green) and GPe population (column iii, red) for two values of the GPe to GPe conductance (scale 0.33; 2.0 in rows C; D respectively). Column ii shows phase vectors of the STN and GPe populations (in green; red, respectively, mean population vectors plotted as thick solid lines and cell vectors as thin transparent lines) reflecting phase locking to the instantaneous GPe phase. STN neurons show a preferred phase of firing but weak beta synchronization, while GPe neurons synchronize more strongly. **Ei**: Population vector length and angle of STN and GPe population (green; red, respectively). **F**: Metrics that characterize bursting in STN neurons: median burst rate, intra-burst firing rate, and coefficient of variation of ISIs across all STN cells. High STN inhibition favors strong bursts (high intra-burst firing rate) and lower inhibition regularization the firing rate (lower CV).

### Increased GPe-STN inhibition shifts STN neurons to a sparse burst firing mode

Increasing the synaptic conductance of the GPe to STN projection shifted the firing mode of STN neurons towards sparse, longer bursts with high intra-burst firing rate set against a lower background firing rate, characterized by a high coefficient of variation of inter-spike intervals (ISIs) (Fig 4.F). Bursting was periodic at low frequencies (~2-5 Hz) but was not synchronized between cells. While STN neurons were weakly entrained to the beta oscillation in the GPe they preferentially fired in an interval leading the GPe by approximately 65 degrees (Fig 4.E). Longer bursts with increased firing rate were promoted by higher availability of voltage-sensitive *Na*^+^ and *Ca*^2+^ channels under more hyperpolarized conditions [39, 43, 52]. This shift towards low-frequency strong bursting was accompanied by increased sub-beta band power, and intra-population synchronization as measured by the vector length (Fig 4, B-C).

**Fig 4.**
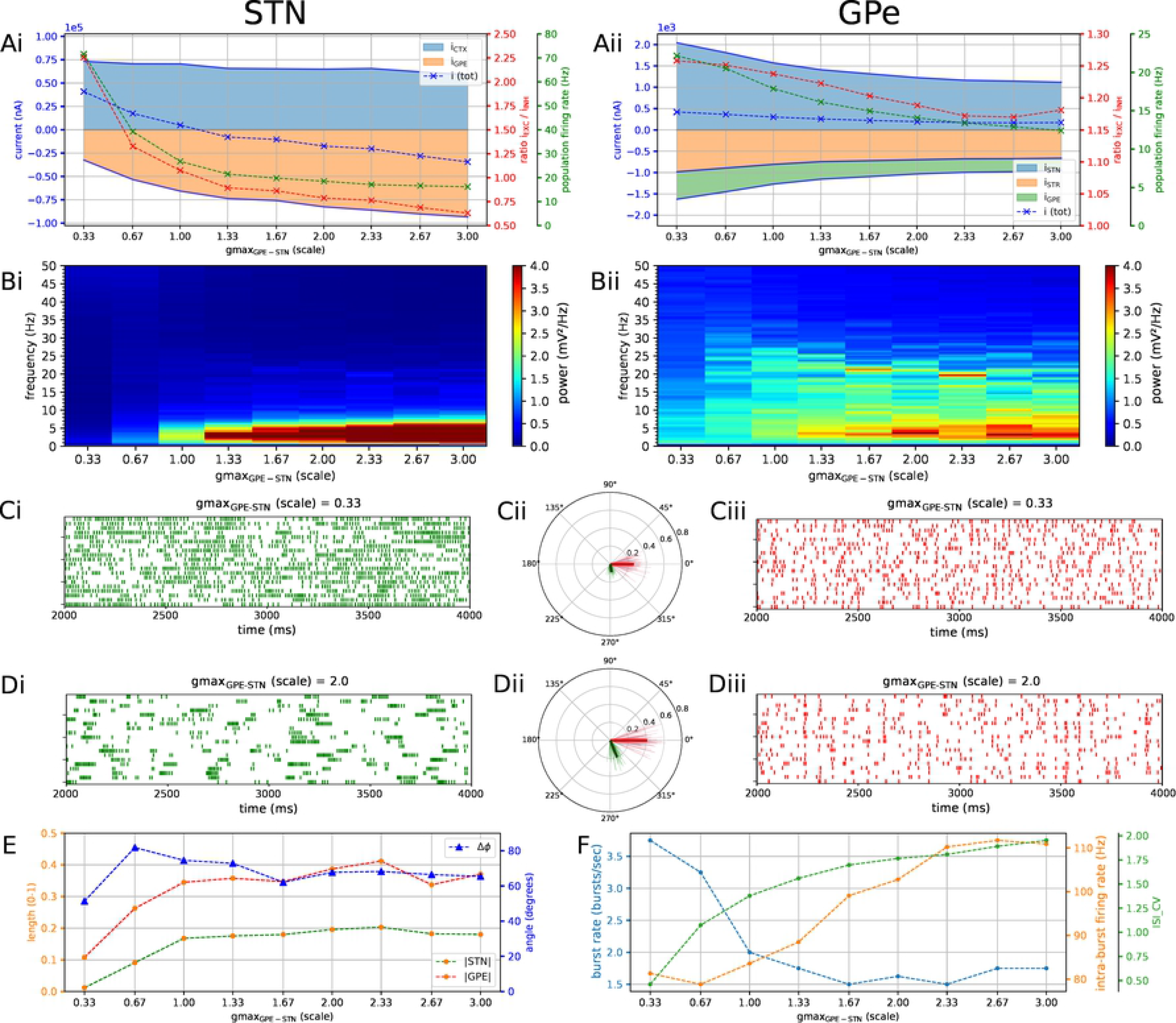
Increasing the level of GPe-STN inhibition shifts STN to a desynchronized low-frequency burst firing mode. Characterization of activity in the autonomous STN-GPe network for increasing values of the GPe to STN synaptic conductance. **A**: Population firing rate (green), E/I ratio (red), and net synaptic current (blue) in STN (Ai) and GPe (Aii). Shaded areas represent estimated total synaptic current from one pre-synaptic population during a simulation. **B**: Mean PSD of the somatic membrane voltages of STN (Bi) and GPe (Bii) neurons. **C-D**: Representative spike trains and phase vectors for STN (column i, green) and GPe population (column iii, red) for two values of the GPe to STN conductance (scale 0.33; 2.0 in rows C; D respectively). Column ii shows phase vectors of the STN and GPe populations (in green; red, respectively, mean population vectors plotted as thick solid lines and cell vectors as thin transparent lines) reflecting phase locking to the instantaneous GPe phase. **E**: Population vector length and angle of STN and GPe population (green; red, respectively). **F**: Metrics that characterize bursting in STN neurons: median burst rate, intra-burst firing rate, and coefficient of variation of ISIs across all STN cells.

### Slow GABA_B_ mediated currents counteract beta patterning of individual STN neurons

With Poisson distributed cortical inputs, GPe neurons were unable to effectively pattern STN neurons even when the GPe neurons exhibited a prominent beta rhythm, despite the increase in GPe to STN conductance. We hypothesized that this was due to the GABA_B_ receptors that provided a slowly decaying baseline level of inhibitory current in our model. This hypothesis was tested by changing the relative strength of the GABA_A_ and GABA_B_ components of the synaptic current while keeping the ratio of excitatory to inhibitory currents in both STN and GPe approximately constant (Fig 5). This was done by increasing the GABA_A_ conductance by a fixed scale factor and scaling down the GABA_B_ conductance by a compensation factor that approximately conserved the total current delivered to STN neurons. The experiment confirmed that the slow nature of the inhibitory current prevented GPe neurons from patterning their targets with brief pauses required for strong entrainment in the 20-30 Hz range. When the proportion of GABA_A_ synapses was increased, with sharp inhibitory post-synaptic currents, and a weak or absent slow GABA_B_ component, the GPe beta rhythm was reliably transmitted to STN neurons and they became completely entrained (Fig. 5.Di).

**Fig 5.**
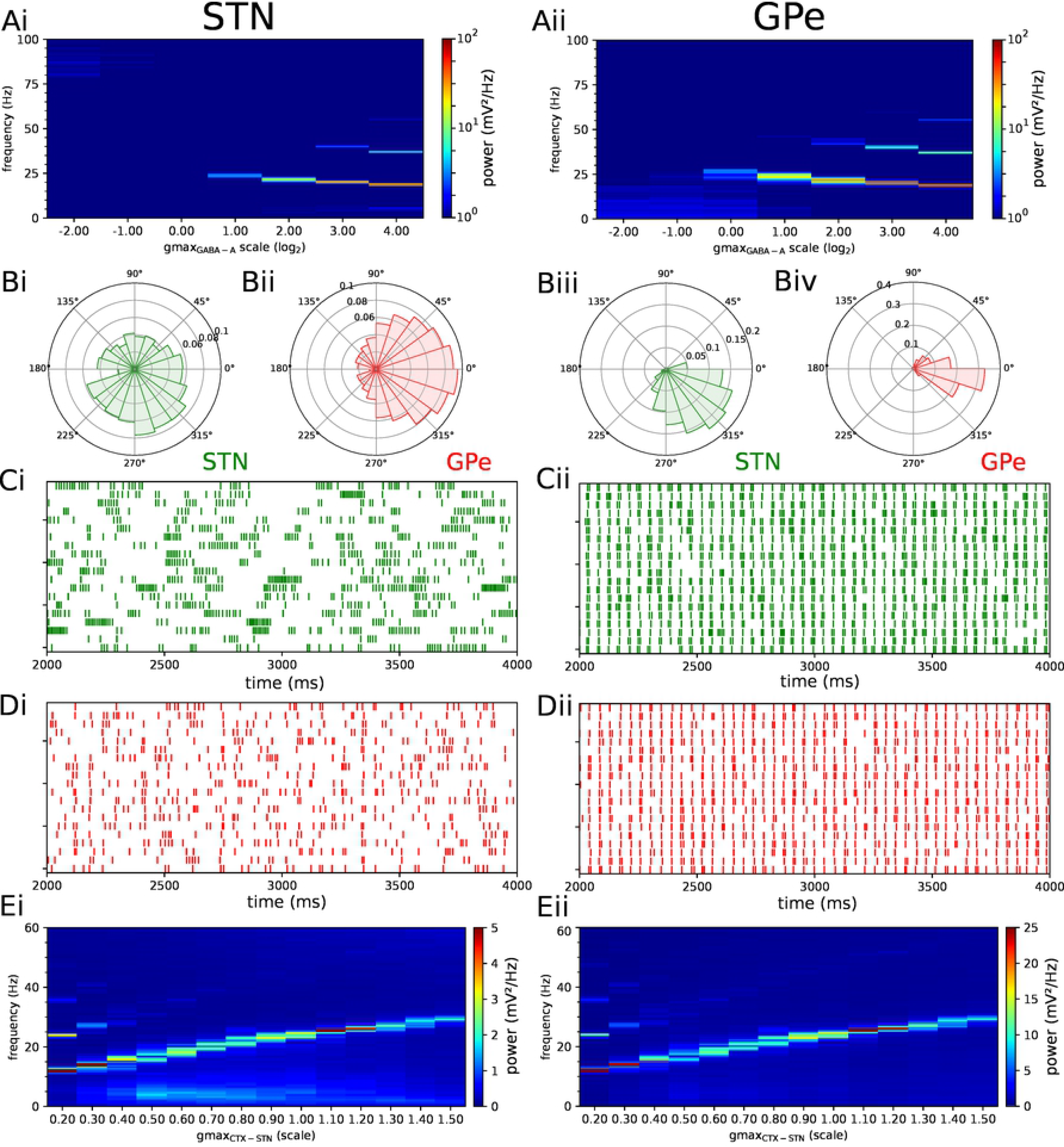
Endogenous oscillations in the STN-GPe network are strengthened by shifting the GPe to STN synaptic current from slow GABA_B_ receptors to fast GABA_A_ receptors. **A**: Reduced GABA_B_R-mediated currents and increased GABA_A_R-mediated currents lead to strengthening of endogenous oscillations. The GABA_B_ conductance of the GPe to STN projection was decreased by 50% and the GABA_A_ conductance was increased progressively (horizontal axis). **B-D**: Effect of shifting the inhibitory current from GABA_B_ to GABA_A_ receptors while keeping the E/I ratio in each population approximately constant. The GABA_A_ conductance was increased by a factor six and the GABA_B_ conductance was adjusted so that the E/I ratio was close to that in the baseline model (left column: baseline model, E/I ratio was 0.89; 0.97 in STN, GPe respectively; right column: scaled conductances, ratio was 0.89; 0.93). **B**: Phase histograms of STN (green) and GPe (red) neurons in baseline model (left; Bi-Bii) and model with scaled conductances (right; Biii-Biv). **C-D**: Representative spike trains of STN (green) and GPe (red) neurons in baseline model (left; Ci-Di) and model with scaled conductances (right; Cii,Dii). **E**: Repeat of the parameter sweep of the CTX to STN conductance in Fig. 2, with scaled GABA conductances. The GABA_B_ conductance of GPe to STN synapses was halved, and the GABA_A_ conductance was doubled. Mean PSD of somatic membrane voltages of STN (Ei) and GPe (Eii) neurons. The oscillation is stronger compared to the baseline model, is reliably transmitted to STN, and the frequency is shifted by the level of cortical excitation.

### STN-GPe network shows resonant properties and phase locks to cortical beta inputs

According to the cortical patterning hypotheses, coherent beta oscillations in the STN and GPe arise due to phase locking of the STN-GPe network to cortical beta-range inputs that are amplified within the loop. To assess the degree of phase locking to such cortical rhythms and its dependence on intrinsic parameters of the STN-GPe network, the network was simulated with cortical inputs modeled as oscillatory beta bursting spike trains (Fig 6). To allow comparison of the behavior of the STN-GPe with exogenous oscillatory inputs with that without oscillatory input shown in Figs. 2-5, the network was simulated for the same range of net excitation-inhibition. This maintained the mean STN firing rate approximately within the experimentally reported range of 17-37 Hz [2, 53] in the dopamine depleted state during cortical activation (compare Fig 2.Aii - Fig. 6.Aii). The STN-GPe network phase locked to beta-band oscillations applied via cortico-STN afferents and the strength of phase locking to a specific frequency was modulated by the excitation-inhibition balance in the same way that it set the spontaneous synchronization frequency (compare to Fig 2). The E/I ratio where the resonance peak occurred corresponded to the E/I ratio where the strength of autonomous oscillation at that frequency was maximal (Fig 6.E). GPe neurons synchronized stronger to the applied beta rhythm compared to STN neurons, which showed a tendency to burst, mirroring the results for spontaneous synchronization in the autonomous STN-GPe network. Analogous to the autonomous loop, when the slow bursting behaviour was reduced by shifting the GPe to STN synaptic current from GABA_B_ to faster GABA_A_ receptors, synchronization and phase locking of both STN and GPe neurons was greatly increased (data not shown).

**Fig 6.**
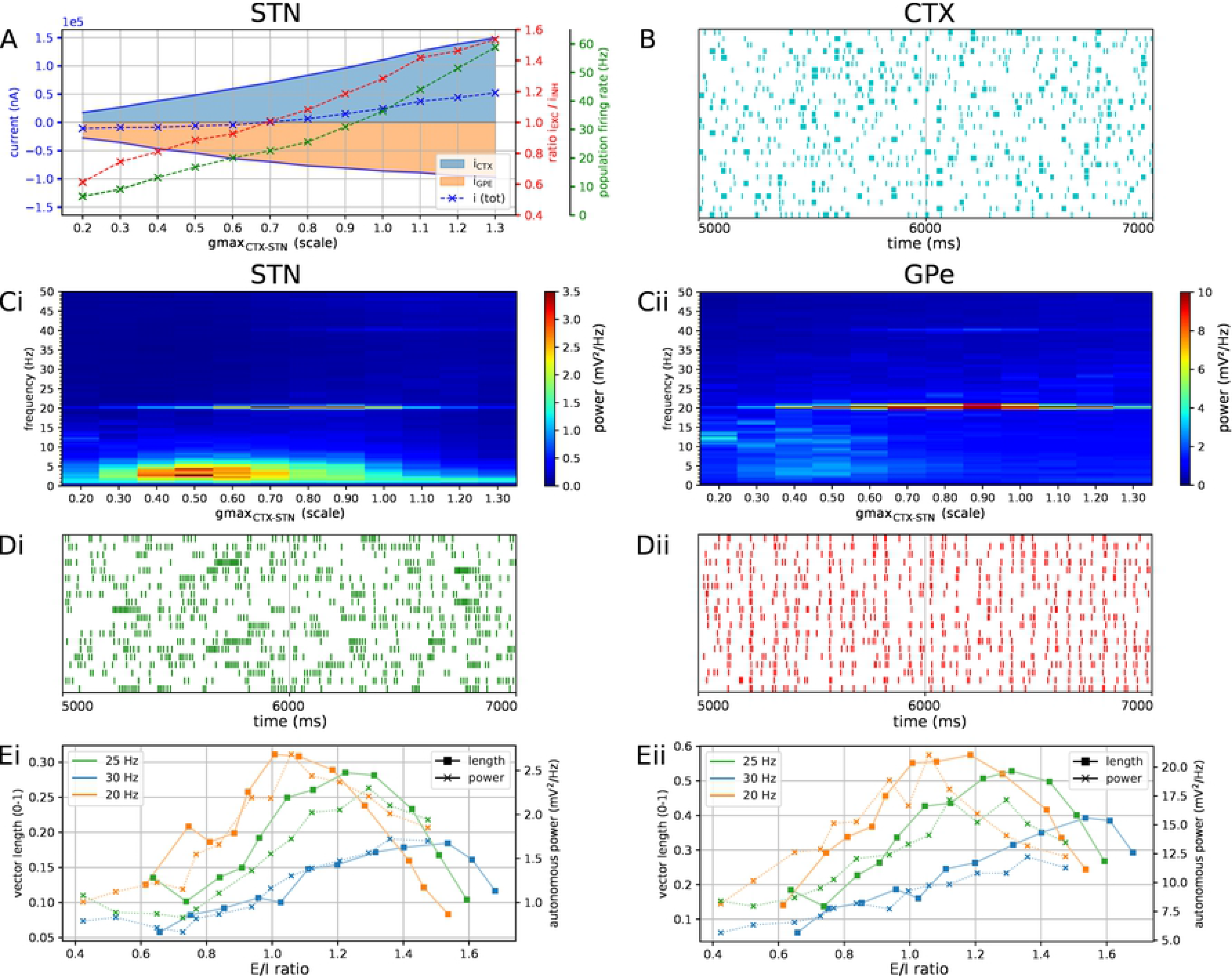
Phase locking of the STN-GPe network to cortical beta-band oscillations at 20, 25, and 30 Hz. Response of the STN-GPe network to oscillatory bursting inputs with different oscillation frequencies applied via cortical afferents. The network model was simulated for each combination of synaptic conductance and bursting frequency. **A**: Population firing rate (green), E/I ratio (red), and net synaptic current (blue) in STN. Results are for network with 20 Hz oscillatory bursting inputs. Shaded areas represent estimated total synaptic current from one pre-synaptic population during a simulation. **B**: Example cortical bursting spike trains with oscillation frequency of 20 Hz. In every cycle of the oscillation 10% of cells were selected at random to fire a burst in phase with the oscillation, with a variation of 1 ms on the onset and spike timings. **C**: Example PSD of the somatic membrane voltages of STN (Ci) and GPe (Cii) across parameter sweep with cortical oscillation frequency set to 20 Hz. **D**: Representative spike trains of STN (Di, green) and GPe neurons (Dii, red) in model with synaptic conductance scale set to 0.7, corresponding to E/I ratio of resonance peak. **E**: Population vector length in network with cortical beta inputs (left axis) and beta-band power in the autonomous network (right axis) for STN (Ei) and GPe (Eii). Autonomous beta-band power was calculated around input frequency in the equivalent network with cortical inputs (bin width = 5 Hz). The x-axis shows the estimated balance of excitatory and inhibitory currents measured from 3 recorded neurons in each population. Maximum phase locking to cortical frequency occurs at approximately the same E/I ratio of maximum autonomous power at the same frequency. Simulations were run with three cortical oscillatory frequencies (20, 25, 30 Hz).

### Preferred phase relationship between cortical and striatal beta inputs

Striatal microcircuits exhibit beta-band oscillations in healthy primates [54] and parkinsonian mice models [23] and have been hypothesized to be part of the pacemaking circuit that generates them. In the previous section, the STN-GPe network was shown to generate weak beta-band oscillations in the absence of exogenous beta inputs (Fig. 2, 3, 4), and to phase lock to cortical beta-band inputs which amplified oscillatory activity (Fig. 6). A potential role of the pallido-striatal loop could be to amplify beta-band oscillations in the STN-GPe network to a more pathological level, as part of a double resonant loop converging on the GPe. A suggested mechanism is that altered striatal activity in PD could shift the phase of firing of the GPe relative to the STN to one that supports STN phase locking through increasing the availability of *Na*^+^ and *Ca*^2+^ channels post-inhibition and pre-excitation [9, 39, 41]. Alternatively, oscillations that have their origin in striatal circuits could be transmitted via the striato-pallidal projection and thus introduced into the STN-GPe network [23, 24]. Of the two loops converging on GPe neurons, inhibitory striatal afferents would be better suited to interrupt ongoing activity and influence the phase compared to excitatory STN afferents. Hence, the MSN to GPe projection could play an important role in patterning neural activity in the STN-GPe network.

Phase vector plots in the previous section show that STN and GPe neurons settle into a particular phase relationship where STN leads GPe by 60-90 degrees which contributed to sustaining beta-band oscillations. Our hypothesis was that inhibitory inputs from the striatum would either disrupt this phase relationship, thereby suppressing beta-band oscillations, or reinforce them depending on the phase of the beta oscillation they arrive. To investigate this hypothesis, surrogate striatal spike trains exhibiting beta bursting activity were generated and the phase with respect to the ongoing oscillation within the STN-GPe network was changed in increments of 45 degrees by varying the onset time of bursts. As MSN-GPe synapses exhibit short-term facilitation, bursts administered through this projection led to an increase in inhibition to the GPe that was greater than the relative increase in spike rate. To compensate for this effect and return the GPe neurons into a more physiological firing rate range, the peak conductance of MSN-GPe synapses was reduced by 60%.

Varying the phase of arrival of striatal bursts revealed that populations connected by an inhibitory projection, i.e. MSN, GPe, and STN maintained a rigid phase relationship with respect to the CTX (Fig. 7: population vectors in green, red, purple formed a rigid frame that rotated relative to the cyan-colored cortical population vector). The local maximum in phase locking occurred when excitatory CTX and inhibitory GPe afferents to STN fired in anti-phase, occurring when the CTX-MSN phase difference was set to 225 degrees (Fig. 7 B,D,Ei). This supports the feedback inhibition hypothesis where cortical patterning is promoted when GPe-STN inhibition is offset in phase relative to cortical excitation in PD [9, 39, 41]. The changing phase relationship of cortical spiking relative to the three other populations also shifted the balance of excitatory and inhibitory currents in STN (Fig. 7.Ai). Maximum phase locking occurred where the STN was maximally inhibited (E/I ratio ≈ 1.1, population firing rate ≈ 21 Hz), whereas minimum phase locking coincided with maximum excitation (E/I ratio ≈ 1.3, population firing rate ≈ 40 Hz). In the GPe this relationship between phase locking strength and firing rate was reversed (Fig. 7.Ai) whereas the relationship with E/I ratio showed no clear trend. The optimal phase relationship of 225 degrees further strengthened phase locking to the applied beta rhythm compared to the situation with only cortical oscillatory inputs. Maximum vector length was increased by a factor of two in both populations compared to the situation with only cortical beta-range inputs, and maximum power at the oscillation frequency was increased by a factor of approximately 2.7 in STN and 5.2 in GPe.

**Fig 7.**
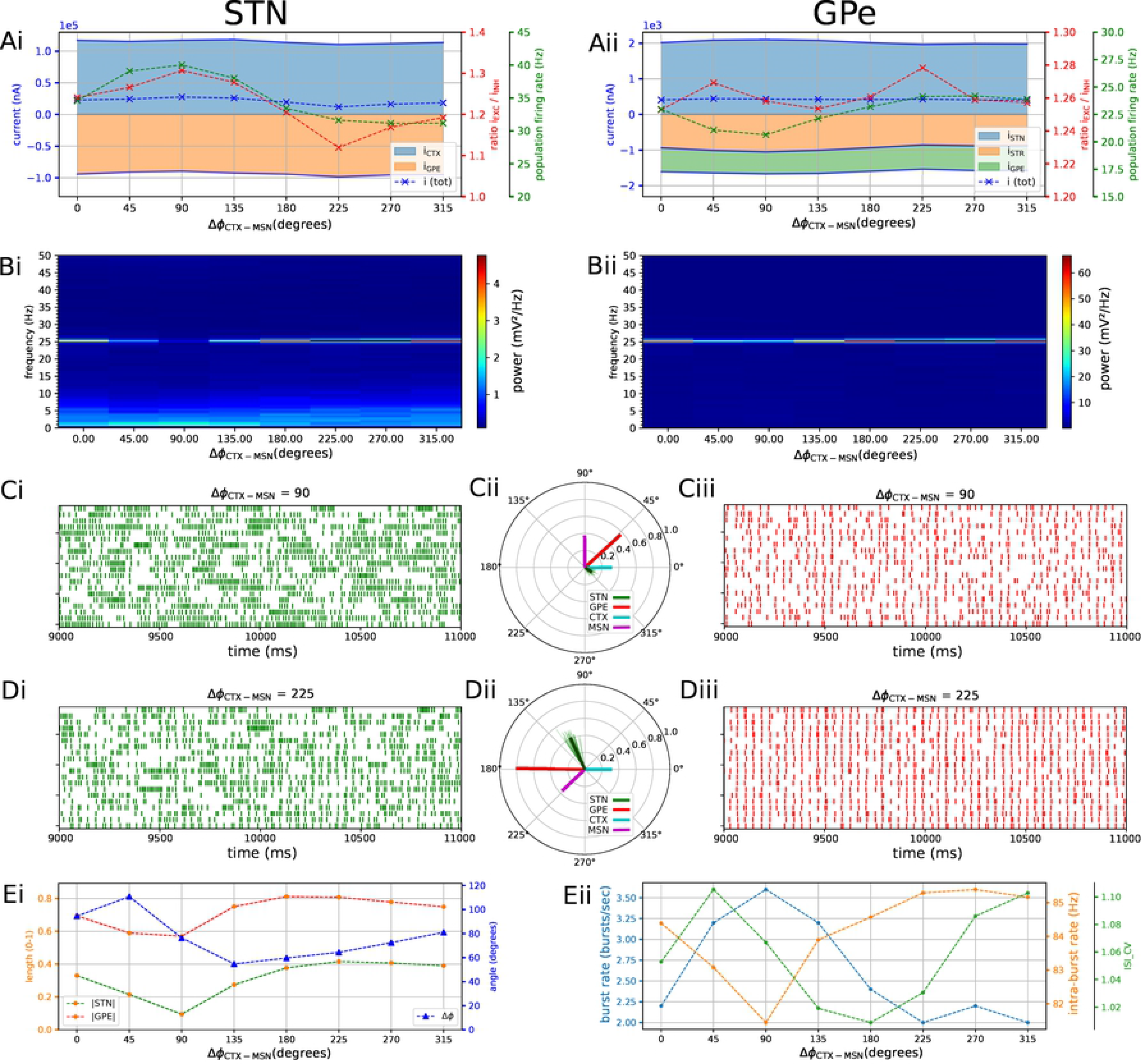
The phase relationship between cortical and striatal beta-band inputs to the STN-GPe network affects the strength of phase-locking by setting the relative timing of excitatory and inhibitory STN afferents. Response of the STN-GPe network to oscillatory bursting inputs applied via both cortico-subthalamic (CTX to STN) and striato-pallidal (MSN to GPe) afferents. The phase difference between cortical and striatal oscillatory spike trains was increased in steps of 45°. **A**: Population firing rate (green), E/I ratio (red), and net synaptic current (blue) in STN (Ai) and GPe (Aii). Shaded areas represent estimated total synaptic current from one pre-synaptic population during a simulation. **B**: Mean PSD of the somatic membrane voltages of STN (Bi) and GPe (Bii) neurons, showing weakening and strengthening of oscillations as relative phases of inputs are rotated. **C-D**: Representative spike trains and phase vectors of STN (column i, green) and GPe population (column iii, red) for CTX-MSN phase difference of 90° (C) and 225° (D). Column ii shows phase vectors of the STN, GPe, CTX, MSN populations (in green; red; blue; purple, respectively; mean population vectors plotted as thick solid lines and cell vectors as thin transparent lines), conditioned on the instantaneous phase of CTX. **E**: Population vector length and angle of STN and GPe population (green; red, respectively). **F**: Metrics that characterize bursting in STN neurons: median burst rate, intra-burst firing rate, and coefficient of variation of ISIs across all STN cells.

### Mechanism of phase locking

The mechanism of phase locking of STN cells is illustrated in Fig. 8. Pooled cortical spike trains (Fig. 8.A-B, green) show how sparse cortical beta bursts (Fig. 6.B) result in distributed synaptic inputs to individual STN neurons that are not tightly phase locked, but have a combined firing rate that is modulated at the beta frequency (Fig. 6.B). While these exogenous cortical inputs had high spike timing variability, STN and GPe spikes became highly structured and tightly locked to the beta oscillation through the feedback inhibition mechanism. The cortical beta modulation is transmitted to the STN and then to the GPe through their excitatory projections (see phase vectors in Fig. 7.Dii). When the inhibitory feedback arrives back in STN this shuts down spiking (Fig. 8.A) and simultaneously primes the cell for the next period of increased cortical excitation by de-inactivating *Ca*^2+^ channels (Fig. 8.C) and *Na*^+^ channels. As the cortical firing rate rises again, synaptic currents (Fig. 8.B) combine with dendritic *Ca*^2+^ currents to overcome any lingering inhibition and cause the next wave of phase-locked STN spikes. The striatal beta inputs further decreased spiking variability of GPe neurons by narrowing their time window of firing through phasic inhibition (purple phase vector in Fig. 7.Dii).

**Fig 8.**
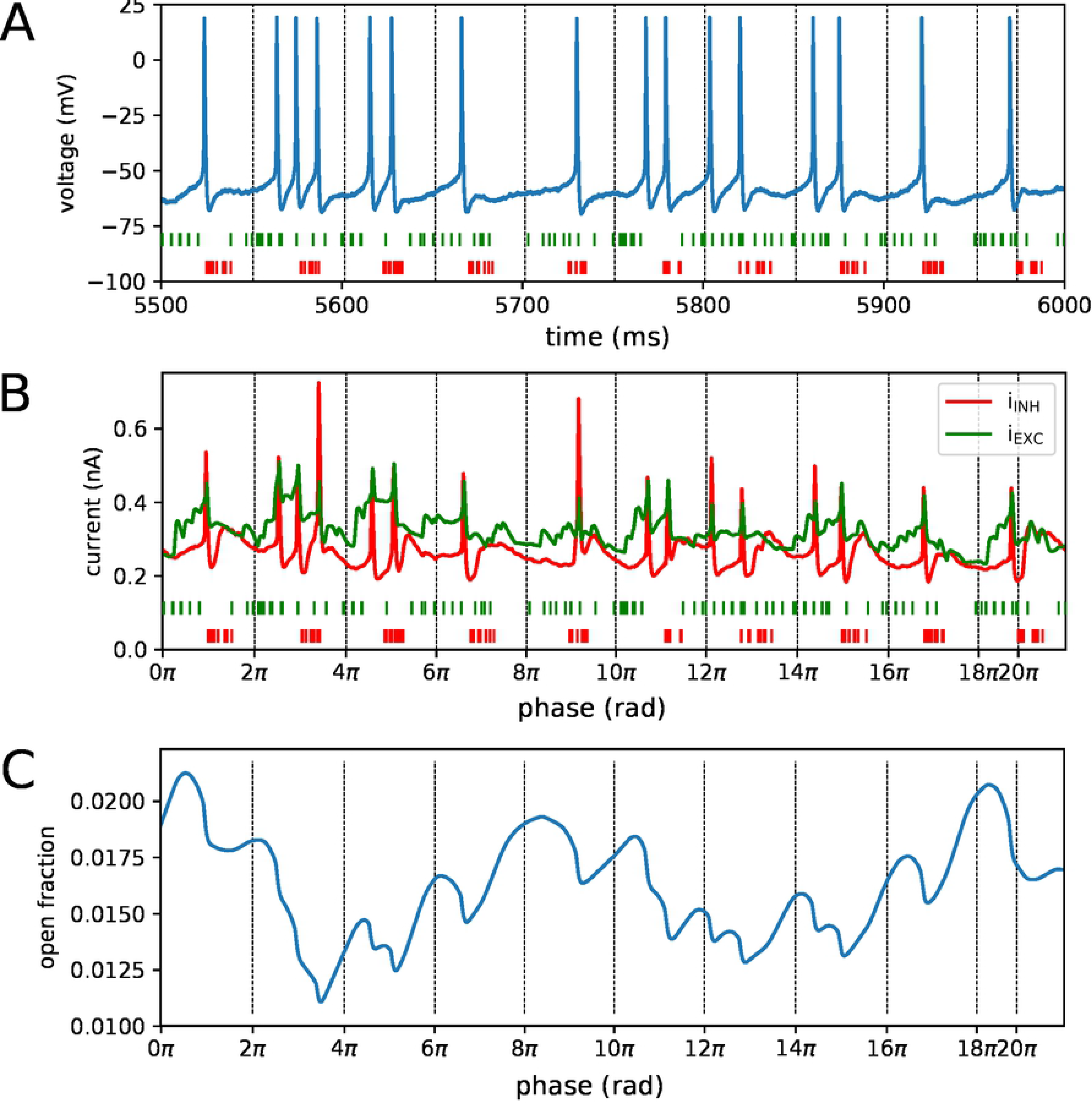
Mechanisms contributing to phase locking of STN cells to cortical beta oscillations. Recordings of synaptic currents and T-type calcium (CaT) channel inactivation from an identified phase-locked STN cell during a simulation with high phase locking (analogous to Fig. 7.D, cortical and striatal beta bursts at 20 Hz with phase difference of 225 degrees). Inactivation variables were recorded from each compartment with CaT ion channels and averaged over all compartments in the cell. Zero-crossings of the instantaneous beta phase are indicated using vertical dotted lines. **A**: Somatic membrane voltage during phase-locked interval. Spike trains from excitatory and inhibitory afferents to the cell were pooled and shown in green and red, respectively. **B**: Total excitatory and inhibitory synaptic current (in green; red, respectively) and pooled spike trains underneath. **C**: Mean CaT channel inactivation across the cell’s dendritic tree. High values correspond to de-inactivation. Transient de-inactivation approximately one half period after an inhibitory barrage engages depolarizing T-type *Ca*^2+^ current and contributes to phase-locked spiking.

## Discussion

A new model of the STN-GPe network is presented that incorporates biophysically detailed cell models and the interaction of intrinsic and synaptic membrane currents with nonuniform subcellular distribution. The model illustrates how phase locking of STN and GPe neurons and exaggerated bursting in STN neurons can arise from the interaction of these currents when their balance and temporal relationships are changed. The model showed an intrinsic susceptibility to beta-band oscillations that was manifest in weak autonomously generated endogenous oscillations and in selective amplification of exogenous beta-band synaptic inputs at the network’s preferred oscillation frequency, for a given excitation-inhibition balance. Varying the phase relationships between oscillatory population spiking patterns increased or suppressed this amplification by modulating spike timing variability and thereby the strength of phase locking. Varying synaptic strengths within the network affected the balance of excitation and inhibition in both STN and GPe neurons and produced a rich set of behaviors, more complex than simple modulation of firing rates. Homeostatic mechanisms mediated by feedback connections and short-term synaptic plasticity dynamics served to stabilize the excitation-inhibition balance in the GPe and reduced the sensitivity of its population firing rate to variations in pre-synaptic rates.

### Oscillatory properties

In the autonomous STN-GPe network, under conditions of Poisson distributed external synaptic inputs, STN neurons exhibited weak synchronization to the endogenous beta rhythm and retained a preferred phase of firing with respect to the GPe oscillations which remained stronger throughout all simulations (Fig. 2-4). The synchronization strength of STN neurons was found to depend on the relative strength of GABA_A_ and GABA_B_ receptors in STN dendrites (Fig. 5), with an increase in the proportion of fast-acting GABA_A_ receptors resulting in an increase in the strength of oscillation. The oscillation frequency of the STN-GPe network was determined by the balance of excitatory and inhibitory currents in the STN. This balance determined the net level of excitatory drive in the loop, shifting the oscillation frequency towards the higher beta range for increased levels of drive (Fig. 2, 6.E). Besides affecting population firing rates and oscillation frequencies, the excitation-inhibition balance also strongly influenced the firing pattern of STN neurons: for a low ratio of excitation to inhibition and sufficiently strong inhibitory currents, STN neurons transitioned to a firing mode characterized by low-frequency tight bursts (high intra-burst firing rate, Fig. 2-4). Low-frequency bursting was periodic at 2-5 Hz but was not synchronized between cells. This shift in firing pattern towards sparse, tight bursting is in correspondence with changes in burst-related measures like intra-burst firing rate and sub-beta band power that are most predictive of akinetic-bradykinetic symptoms in humans [11] and monkeys [10], although sub-beta band occurs at a higher frequency in monkeys. The firing rate and pattern of GPe neurons was less sensitive than STN neurons to variations in its excitatory or inhibitory drive as it was under negative feedback control by homeostatic mechanisms that operated in synergy to stabilize its E/I ratio. However, GPe neurons did synchronize more strongly under conditions of low excitatory drive from the STN enabling them to act more autonomously and synchronize through inhibitory collaterals.

When beta-band spiking inputs were applied via cortico-STN afferents, the STN-GPe network phase locked to the beta rhythm and input frequencies near the autonomous oscillation frequency for a E/I ratio were preferentially amplified, reflected in increased phase locking and oscillatory power of the somatic membrane voltage (Fig. 6.C,E). This is supportive of experimental observations that oscillatory activity in STN is contingent on cortical oscillations [38], likely transmitted though the hyperdirect pathway [12]. Phase locking and oscillation power were further strengthened by addition of striatal oscillatory inputs with a particular phase relationship to cortical oscillatory inputs (Fig. 7). The maximum in phase locking occurred when cortical and GPe afferents to STN were pushed to anti-phase by the timing of striatal inputs. This anti-phase relationship made GPe inhibition more effective at facilitating bursting of STN neurons by hyperpolarizing them, thereby de-inactivating *Ca*^2+^ and *Na*^+^ channels and priming them for a response to the next wave of cortical excitation (Fig. 7.A: maximum phase locking coincided with maximal phase-dependent inhibition). On the other hand, phase alignment of cortical and GPe neurons, corresponding to coincident firing, desynchronized STN neurons (Fig. 7.C). This is in agreement with recent experimental findings showing co-stimulation of GABAergic and glutamergic STN afferents disperses STN spiking and has a desynchronizing effect [55]. Overall, the simulation results are consistent with the hypothesis of cortical patterning through feedback inhibition, where GPe inhibition arriving in anti-phase to cortical excitation promotes phase locking of STN neurons to beta-band cortical inputs [39].

### Relation to other models of oscillations

The mechanism by which activity in the STN and GPe oscillates by means of alternating phases of excitation and inhibition in a delayed negative feedback loop has been described in previous models [19, 21, 48]. The mechanisms of oscillation in our model of the STN-GPe network is consistent with this, but further illustrates the dual role of GPe inhibition of STN neurons in shutting down ongoing activity and priming for the next wave of excitation (Fig. 8). Moreover, in our model endogenously generated oscillations were weak in the autonomous STN-GPe network in all simulation except when fast-acting inhibitory currents in STN dominated strongly over slow ones. Spiking variability was sufficiently decreased to cause a strongly synchronized beta-band oscillation to emerge only when exogenous beta-range inputs were applied. (Fig. 6, 7).

The role of such patterning input could be fulfilled either by an extrinsic oscillator or by reverberation of oscillations in connected feedback loops, e.g. pallido-striatal circuits [24], recurrent striatal loops [23], or the thalamocortical backprojection [25, 28, 56]. Our results, therefore, support the role of resonance with other basal ganglia loops in generating exaggerated synchrony. Our model showed that the preferred oscillation frequency could be modulated by varying the ratio of excitation to inhibition in STN and GPe (Fig. 2B, 6.E) and that the ranges of strong oscillations and low-frequency bursting overlapped (Fig. 6.C). Mean field models have shown that the oscillation frequency is strongly modulated by transmission delays and membrane time constants [21, 57], while synaptic strengths have a weaker effect on the oscillation frequency with sensitivity comparable to that in our model [21, 22]. However, in a model incorporating active ion channels that contribute to synaptic integration, synaptic strength and effective membrane time constant cannot be strictly separated since the membrane charging speed is affected by ion channels that are transiently activated under the influence of synaptic stimulation.

We did not investigate transient fluctuations in beta power which have been observed in health and disease, and are linked to motor performance [58, 59] although such fluctuations were observed in spectrograms (data not shown). These fluctuations have been hypothesized to be cortical in origin, originating in basal ganglia structures, or arising due to the interaction of the two ([60]). In our model, low-frequency fluctuations in beta-band power could arise from calcium-activated potassium currents triggered by phases of high activity due to short-term plasticity dynamics as shown in other models [30]. Under the full-loop resonance hypothesis low-frequency ‘waxing and waning’ of beta activity could arise from modulations in beta power in any node of the BGTC network, either through intrinsic dynamics or competition between extrinsic inputs.

### Model complexity and limitations

One of the main advantages of using a biophysically detailed model that incorporates the level of detail presented here is that afferents from each pre-synaptic population can target a specific region of the dendritic tree (Table 3, 5), leading to differing synaptic integration properties. Specifically in the light of pathological synchronization this is important given that neuronal phase response curves, used to quantify the tendency of neurons to synchronize to their inputs, differ when stimuli are applied to different subcellular regions in the STN and GPe [61, 62]. Hence, a model that incorporates a full complement of ion channel and the synapse groups that interact with them may be expected to yield a more realistic picture of how synchronization arises in the network. In future studies, this can also contribute to a better understanding of neuronal currents contributing to the local field potential in synchronized and asynchronous states, as synaptic and ionic transmembrane currents combine to form the extracellular currents that underpin this signal [63].

A second advantage of detailed models is that parameters have a clear relationship to the underlying biophysical system and are more meaningful compared to the case where parameters are lumped, as in single-compartment conductance-based models, or abstracted as in mean-field or generalized integrate-and-fire models. This allows for direct translation of experimental findings to parameter variations in the model. On the other hand, detailed cell models are more sensitive to correct estimation of these parameters which is limited by measurements performed for the purpose of model fitting as well as the fitting procedure itself. In this model, downregulation of HCN channel currents under dopamine depletion was modeled as a decrease in its peak conductance. However, dopamine is known to interact with several ion channels that are involved in linearizing the current-firing rate curve and regularizing autonomous pacemaking of STN neurons [64–66] which are not all included in the STN cell model used here [43]. Recent evidence suggests that this loss of autonomous spiking is a necessary condition for the exaggerated cortical patterning of STN that is related to motor dysfunction [67]. Better characterization of the ion channels involved in pacemaking and their response to dopamine depletion will enable the systematic exploration of their contribution to STN response properties and pathological firing patterns.

In our network model the main sources of firing rate variability were randomness in the input spiking patterns, the presence of surrogate Poisson spike sources in STN and GPe, membrane noise, and randomness in connection patterns and the position of synapses. However these factors do not capture the full biological variability in morpho-electric cell types, synaptic strength distributions, and resulting firing patterns in each population. Indeed, in the GPe two distinct populations have been identified based on their molecular profile and axonal connectivity [41]. Of these we only modeled the prototypic population, projecting back to STN and preferentially firing in anti-phase to it. Moreover, the GPe cell model used was only one representative candidate out of a large set of models with varying ion channel expression and morphology that matched a corresponding database of electrophysiological recordings [50]. Likewise, the STN model represents a stereotypic characterization and does not capture variability in firing properties and receptor expression. In particular, STN neurons in vivo are known to have variable expression of GABA_B_ receptors [44] which cause strong hyperpolarization responses and longer pauses in some but not all STN neurons [52] and a strong rebound burst response [44] in a subset of these. A model that takes into account the biological variability in GABA_B_ expression and that of channels underlying the rebound response may reveal a wider range of responses to increased inhibition among STN neurons. In such a model, beta rhythms could be transmitted to a subset of STN neurons whereas others would show longer pauses with stronger rebound bursts. Moreover, the GABA_B_ synapse model used does not fully account for activation of extrasynaptic GABA_B_R due to GABA spillover [44] which is mediated by tonic high-frequency *and* coincident firing of afferents [40]. A model where multiple GABAergic synapses act on a shared pool of extrasynaptic GABA_B_R might increase the importance of synchronized pre-synaptic activity in switching STN neurons to a burst-firing mode.

To conclude, biophysically detailed models offer new ways to study factors contributing to the development of synchrony. Such models provide a means to investigate the relative contributions of physiological mechanisms to the development of synchrony while controlling other factors in a manner that is not possible in vivo. Despite the relatively high-level of detail of the model, various factors contributing to biological variability and shaping firing neuronal firing patterns were omitted due to the model complexity.

## Conclusion

In summary, a biophysically detailed model of the parkinsonian STN-GPe network is presented which showed an intrinsic susceptibility to beta-band oscillations. However oscillations in the autonomous STN-GPe network were too weak to support a pacemaker role that is the sole origin of beta-band oscillations in the wider BGTC network in Parkinson’s disease. In particular in the STN, autonomous beta-band oscillations and phase locking of individual units were weak unless slower GABA_B_-mediated currents were reduced. Beta-band oscillations were considerably amplified by a relatively sparse cortical beta input, and were further amplified by striatal beta inputs that promoted anti-phase firing of cortex and GPe. These results support the cortical patterning and network resonance hypothesis and point to a role for the pallido-striatal feedback loop in amplifying beta oscillations.

## Methods

### Model architecture

The network model of the STN-GPe network consisted of four populations (Fig. 1): the STN and GPe, modeled as multi-compartmental conductance-based models, and their cortical and striatal inputs, modeled as Poisson or bursting spike generators.

Population sizes were chosen to preserve the decrease in population sizes and convergence of projections along the indirect and hyperdirect pathways in the basal ganglia. The STN and GPe populations consisted of 50 and 100 multi-compartmental cells, respectively, to approximate the ratio of roughly 13,000 STN cells to 30,000 GPe prototypic cells [68, 69] unilaterally in the rat. As a source of synaptic noise, an additional 10% of the cells in the STN and GPe populations were modeled as Poisson spike generators firing at a mean rate equal to the experimentally reported rate for the modeled state.

The cortical and striatal populations consisted of 1,000 and 2,000 cells, respectively, modeled as spike generators. These numbers were chosen to have 20 independent pre-synaptic spike generators per post-synaptic cell to model convergence along the hyperdirect CTX-STN and indirect GPe-MSN projection. For the GPe-MSN projection, convergence from MSN to GPe, ignoring subpopulations, is approximately 2,800,000 MSN cells to 46,000 GPe cells [70] resulting in a convergence factor of approximately 60. Assuming that convergence is similar between indirect pathway MSN and GPe prototypic neurons, our number is an underestimation by a factor three. Because MSN cells in our model spike independently and since the number of synapses per cell was lower than in reality, this was considered acceptable.

Stochastic connectivity profiles for the connections illustrated in Fig. 1 were generated by randomly selecting a fixed number of afferents from the pre-synaptic population for each post-synaptic cell. The ratios of number of afferents from each source population were determined, where possible, based on the reported number of synaptic boutons per afferent type and the number of contacts per axon. Each multi-synaptic contact was represented by a single synapse to reduce the number of simulated synapses to a more tractable number.

### Conductance-based models

In the conductance-based formalism, the membrane potential *v*_*j*_ (mV) in each compartment *j* of a multi-compartmental cable model is governed by:

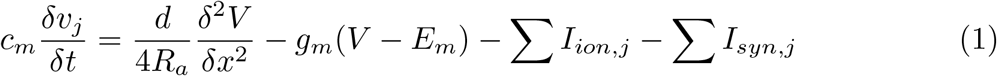

where x (cm) is the position along the cable, *c*_*m*_ (*μF/cm*^2^) is the specific membrane capacitance, *d* (cm) is the cable diameter, *R*_*a*_ is the specific axial resistance (Ω*cm*), *g*_*m*_ (*S/cm*^2^) is the passive membrane conductance, *E*_*m*_ (mV) the leakage reversal potential, *I*_*ion,j*_ (*mA/cm*^2^) are the ionic currents flowing across the membrane of compartment *j*, and *I*_*syn,j*_ (*mA/cm*^2^) are the synaptic currents at synapses placed in the compartment. Each ionic current is governed by an equation of the form:

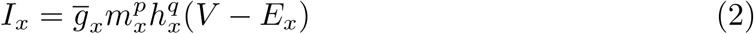

where 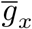 is the maximum conductance of the channel (*S/cm*^2^), *E*_*x*_ is the reversal potential (mV), and *m*_*x*_ and *h*_*x*_ the open fractions of the activation and inactivation gates. The dynamics of the activation and inactivation gates *m* and *h* are governed by

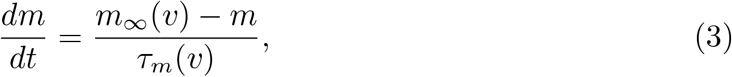

with *m*_∞_(*v*) and *τ*_*m*_(*v*) representing the voltage-dependent steady state value and time constant of the gate. For some currents the gating dynamics are described in terms of the opening and closing rates *α*_*m*_ and *β*_*m*_ related through 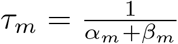, 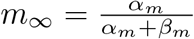:

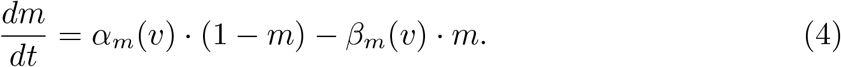

Reversal potentials are assumed constant unless otherwise noted. The reversal potential for Ca^2+^ currents was calculated using the Nernst equation from the intra- and extracellular ion concentrations:

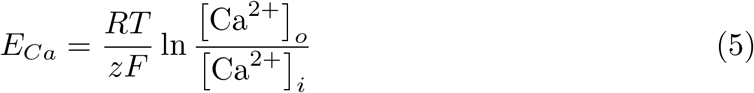

where T is the temperature in Kelvin, R is the universal gas constant, F is the Faraday constant, and z is the valence of the Ca ion (+2). Intracellular calcium buffering in a sub-membrane shell is modeled as:

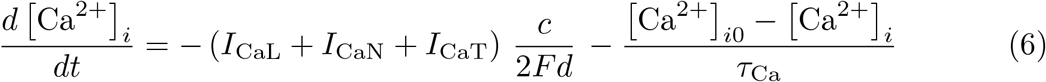

where *c* is a unit conversion constant, d is the thickness of the sub-membrane shell, and *τ*_Ca_ is the time constant of decay.

Synapses were modeled by a dual exponential profile with rise and decay times *τ*_*rise*_ and *τ*_*decay*_ modulated by the fraction of synaptic resources in the active state which was governed by Tsodyks-Markram dynamics [71]:

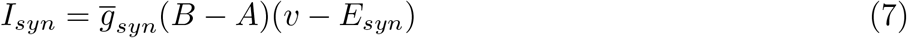

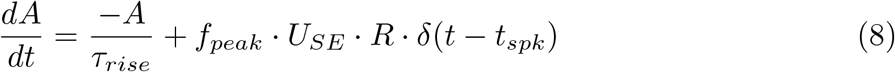

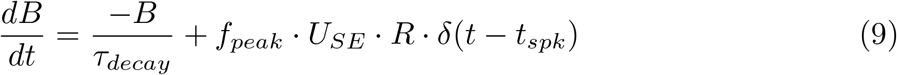

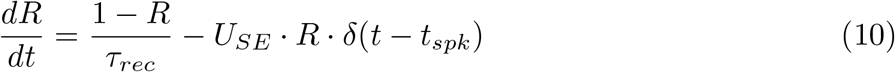

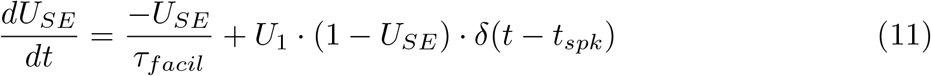

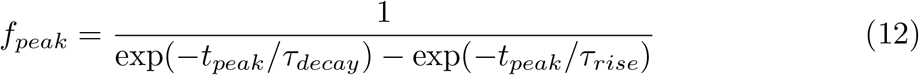

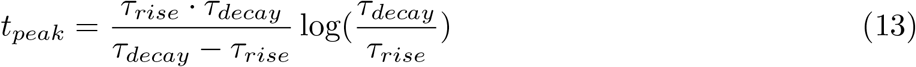

where, 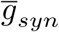 is the peak synaptic conductance, B-A represents the synaptic gating variable, *f*_*peak*_ is a normalization factor so that B-A reaches its maximum at time *t*_*peak*_ after the time of spike arrival *t*_*spk*_, R is the fraction of vesicles available for release, *U*_*SE*_ is the release probability, and *τ*_*rec*_ and *τ*_*facil*_ are the time constants for recovery from short-term depression and facilitation, respectively. The synaptic reversal potentials *E*_*syn*_ were 0 mV for AMPA and NMDA, −80 mV for GABA_A_, and −95 mV for GABA_B_.

For NMDA synapses there is an additional voltage-dependent gating variable representing magnesium block [72]:

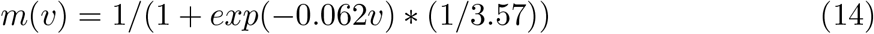

The metabotropoc GABA_B_ receptor-mediated current was modeled as an intracellular signaling cascade based on the model by Destexhe and Sejnowski [73]. The equations describing G-protein activation and the synaptic current were retained, but the bound receptor fraction including the effects of desensitization was represented by the fraction of resources in the active state in the Tsodyks-Markram scheme (B-A). The equation governing the G-protein production rate thus became

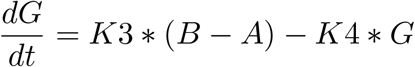

where G is the G-protein concentration, and K3 and K4 are the rates of G-protein production and decay, respectively.

### STN cell model

STN neurons were modeled using the rat subthalamic projection neuron model by Gillies and Willshaw [43] (ModelDB accession number 74298). The neuron morphology is based on quantitative characterization of the dendritic trees of STN neurons in vitro. The model includes ten intrinsic ionic currents (Table 2):

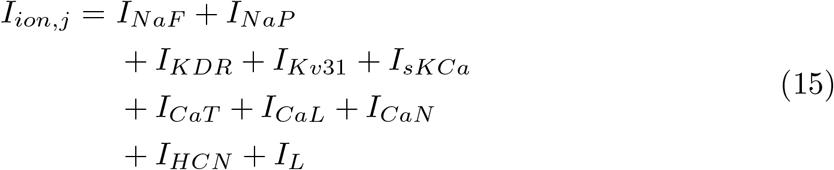

where *I*_*NaF*_ and *I*_*NaP*_ are the transient fast-acting and persistent sodium current, *I*_*KDR*_, *I*_*Kv*31_, and *I*_*sKCa*_ the delayed rectifier, fast rectifier and calcium-activated potassium current, *I*_*CaT*_, *I*_*CaL*_ and *I*_*CaN*_ the low-voltage-activated T-type, high-voltage-activated L-type, and high-voltage-activated N-type calcium currents, *I*_*HCN*_ the hyperpolarization-activated cyclic nucleotide (HCN) current, and *I*_*L*_ the leak current. The equations governing the dynamics of the gating variables are listed in Table 2. The channel density distributions are described extensively in ref. [43]. As a source of noise, a current that followed a Gaussian amplitude distribution with mean zero and standard deviation 0.1 was added to the somatic compartment.

**Table 1.** Experimentally reported connection parameters used to calibrate the model.

| target | source | afferent<br>neurons | synaptic<br>contacts | subcellular<br>targets | short-term<br>plasticity | delay | effect DD |
| --- | --- | --- | --- | --- | --- | --- | --- |
| STN | (all) | 300 [90] |  | N.A. | N.A. | N.A. | N.A. |
|  | CTX |  |  | distal [15, 45, 91],<br>proximal [15] | depression [92] | 5.9 ms [53] | weakened [46, 47] |
|  | GPe | 57 [93] | 883 [94] | proximal [95] | depression [93] | 4 ms [96] | strengthened [92]<br>prolonged decay [74] |
| GPe | GPe |  |  | proximal,somatic [49, 97] | depression [82] |  | strengthened [82] |
|  | STN | 135 [98] |  | dendritic,distal [99] | facilitation,<br>depression [51] | 2 ms [100] | strengthened [78] |
|  | MSN |  | 10622 [98] | dendritic, distal [49] | facilitation [82] | 5 ms [100] |  |

**Table 2.**
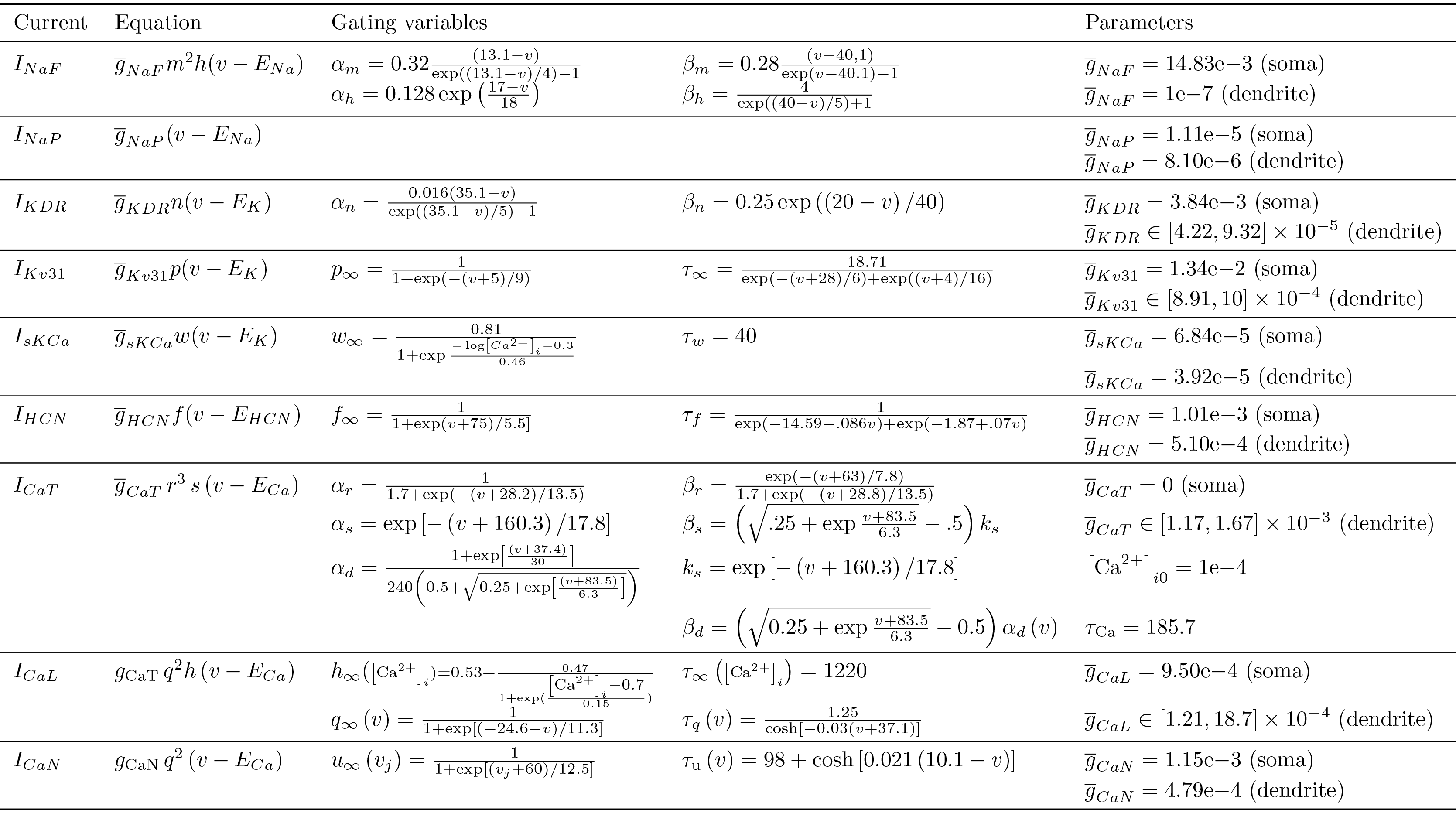
STN model intrinsic current equations from Gillies and Willshaw [43].

The synaptic currents included an excitatory glutamergic input from cortex, acting through AMPA and NMDA receptors, and an inhibitory GABAergic input from the GPe, acting through GABA_A_ and GABA_B_ receptors (Table 3):

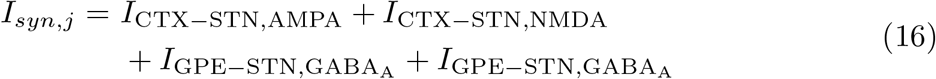

**Table 3.**
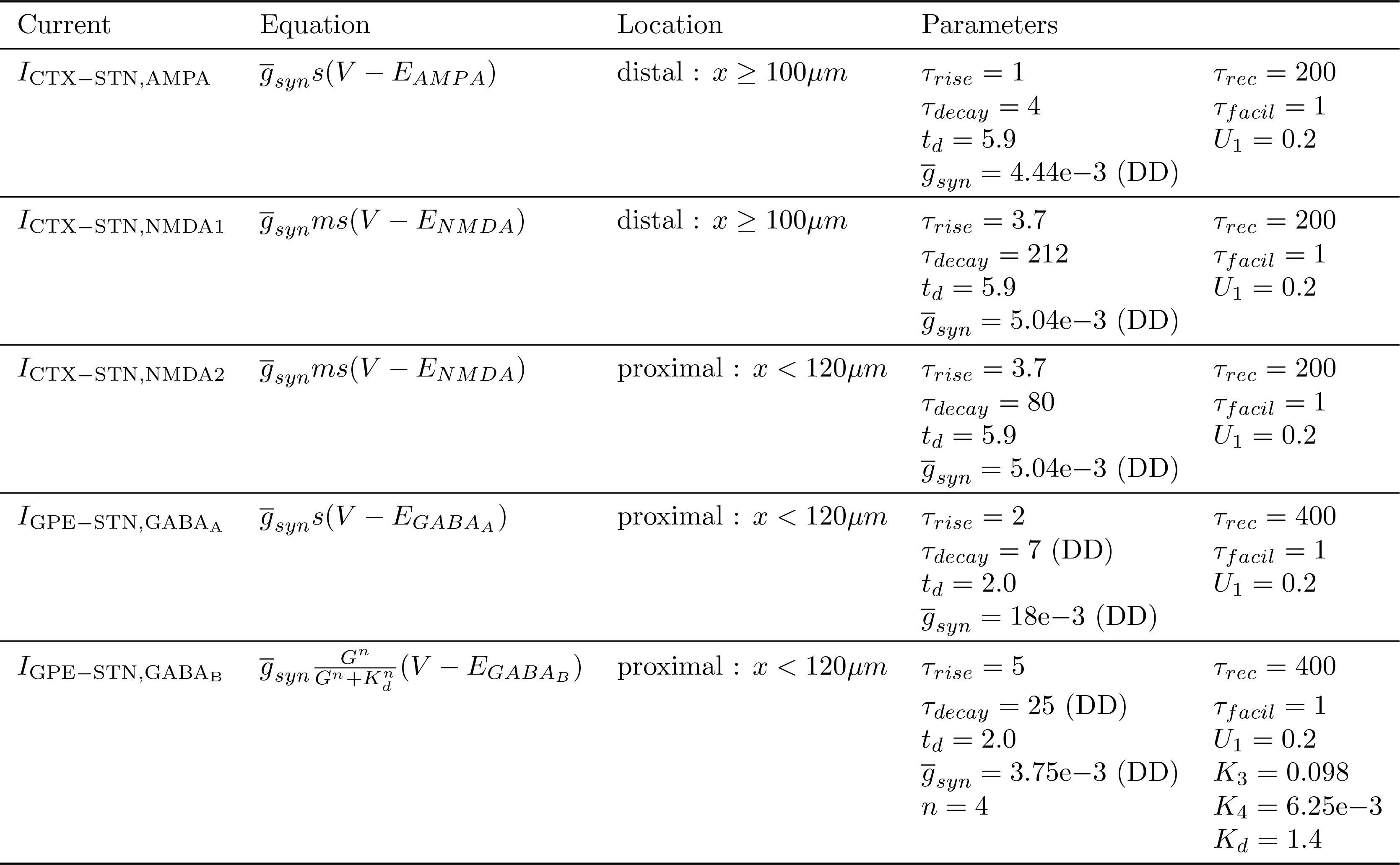
STN model synaptic current equations.

In the control condition STN neurons had 20 excitatory cortical afferents and 8 inhibitory pallidal afferents. Excitatory glutamergic synapses were modeled as conductance-based synapses with Tsodyks-Markram dynamics [71]. For each cortical afferent one model synapse was placed distally in the dendritic tree and one model synapse was placed proximally near the soma. The distal synapses had both an AMPA and slower NMDA conductance component with common short-term plasticity dynamics but separate conductance variables. The latter represented slower-kinetics NMDA receptors with majority NR2B and NR2D subunits that have dendritic punctual expression [15]. The proximal synapses had only an NMDA component and represented NMDA receptors with NR2A subunits that have faster kinetics [15]. The parameter values for the conductance-based synapses are listed in Table 3.

### GPe cell model

GPe neurons were modeled using the baseline rat GPe neuron model by Günay et al. [50] (ModelDB accession number 114639). The model is based on a reconstructed morphology from the adult rat and contains nine types of ion channels with varying densities in the soma, dendrite, and axon initial segment:

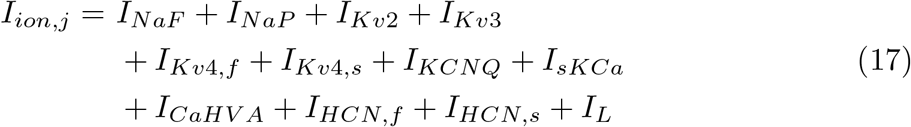

where *I*_*NaF*_ and *I*_*NaP*_ are the transient fast-acting and persistent sodium current, *I*_*Kv*2_ and *I*_*Kv*3_ the slow and fast delayed rectifier potassium current, *I*_*Kv*4*f*_ and *I*_*Kv*4*s*_ the fast and slow component of the A-type, transient potassium current, *I*_*KCNQ*_ the M-type potassium current, *I*_*sKCa*_ the calcium-dependent potassium current, *I*_*CaHV A*_ the high-threshold, noninactivating calcium current (reflecting a mixture of L, N, and P/Q-type calcium channel types), and *I*_*HCN,f*_ and *I*_*HCN,s*_ the fast and slow component of the HCN channel. The equations governing the dynamics of the gating variables are listed in Table 4. The channel density distributions are those described in ref. [50] for model t9842. As a source of noise, a current that followed a Gaussian amplitude distribution with mean zero and standard deviation 0.0075 was added to the somatic compartment. This number was chosen to result in membrane voltage noise of similar amplitude the STN cell model, given the lower somatic input resistance of the latter model.

**Table 4.** GPe model intrinsic current equations from Günay et al. [50].

| Current | Equation | Gate | $m_0$ | $\theta_{m\infty}$ | $\sigma_{m\infty}$ | $\tau_0$ | $\tau_1$ | $\theta_{m\tau}$ | $\sigma_{m0}$ | $\sigma_{m1}$ | Additional parameters |
| --- | --- | --- | --- | --- | --- | --- | --- | --- | --- | --- | --- |
| $I_{NaF}$ | $\bar{g}_{NaF}m^3hs(v - E_{Na})$ | m | 0 | -39 | 5 | 0.028 | 0.028 | N/A | N/A | N/A | $\bar{g}_{NaF} = 0.035$ (soma) |
| | | h | 0 | -48 | -2.8 | 0.025 | 4 | -43 | 10 | -5 | $\bar{g}_{NaF} = 0.035$ (dendrite) |
| | | s | 0.15 | -40 | -5.4 | 10 | 1000 | -40 | 18.3 | -10 | $\bar{g}_{NaF} = 0.5$ (axon) |
| $I_{NaP}$ | $\bar{g}_{NaP}m^3hs(v - E_{Na})$ | m | 0 | -57.7 | 5.7 | 0.03 | 0.146 | -42.6 | 14.4 | -14.4 | $\bar{g}_{NaP} = 10.15e-3$ (soma) |
| | | h | 0.154 | -57 | -4 | 10 | 17 | -34 | 26 | -31.9 | $\bar{g}_{NaP} = 10.15e-3$ (dendrite) |
| | | s | 0 | -10 | -4.9 | N/A | N/A | N/A | N/A | N/A | $\bar{g}_{NaP} = 4e-3$ (axon) |
| $I_{Kv2}$ | $\bar{g}_{Kv2}m^4h(v - E_K)$ | m | 0 | -33.2 | 9.1 | 0.1 | 30 | -33.2 | 21.7 | -13.9 | $\bar{g}_{Kv2} = 0.1e-3$ (soma, dendrite) |
| | | h | 0.2 | -20 | -10 | 3400 | 3400 | N/A | N/A | N/A | $\bar{g}_{Kv2} = 64e-3$ (axon) |
| $I_{Kv3}$ | $\bar{g}_{Kv3}m^4h(v - E_K)$ | m | 0 | -26 | 7.8 | 0.1 | 14 | -26 | 13 | -12 | $\bar{g}_{Kv3} = 1e-3$ (soma, dendrite) |
| | | h | 0.6 | -20 | -10 | 7 | 33 | 0 | 10 | -10 | $\bar{g}_{Kv3} = 128e-3$ (axon) |
| $I_{Kv4,f}$ | $\bar{g}_{Kv4,f}m^4h(v - E_K)$ | m | 0 | -49 | 12.5 | 0.25 | 7 | -49 | 29 | -29 | $\bar{g}_{Kv4,f} = 2e-3$ (soma) |
| | | h | 0 | -83 | -10 | 7 | 21 | -83 | 10 | -10 | $\bar{g}_{Kv4,f} = 4e-3$ (dendrite) |
| | | | | | | | | | | | $\bar{g}_{Kv4,f} = 160e-3$ (axon) |
| $I_{Kv4,s}$ | $\bar{g}_{Kv4,s}m^4h(v - E_K)$ | m | 0 | -49 | 12.5 | 0.25 | 7 | -49 | 29 | -29 | $\bar{g}_{Kv4,s} = 3e-3$ (soma) |
| | | h | 0 | -83 | -10 | 50 | 121 | -83 | 10 | -10 | $\bar{g}_{Kv4,s} = 6e-3$ (dendrite) |
| | | | | | | | | | | | $\bar{g}_{Kv4,s} = 240e-3$ (axon) |
| $I_{KCNQ}$ | $\bar{g}_{KCNQ}m^4h(v - E_K)$ | m | 0 | -61 | 19.5 | 6.7 | 100 | -61 | 35 | -25 | $\bar{g}_{KCNQ} = 20e-5$ (soma, dendrite) |
| | | | | | | | | | | | $\bar{g}_{KCNQ} = 4e-5$ (axon) |
| $I_{CaHVA}$ | $\bar{g}_{CaHVA}m(v - E_{Ca})$ | m | 0 | -20 | 7 | 0.2 | 0.2 | -20 | N/A | N/A | $\bar{g}_{CaHVA} = 3e-5$ (soma, thick dendrites) |
| | | | | | | | | | | | $\bar{g}_{CaHVA} = 4.5e-5$ (medium dendrites) |
| | | | | | | | | | | | $\bar{g}_{CaHVA} = 9e-5$ (thin dendrites) |
| | | | | | | | | | | | $[Ca^{2+}]_{i0} = 5e-5$<br>$\tau_{Ca} = 1$ |
| $I_{HCN,f}$ | $\bar{g}_{HCN,f}m(v - E_h)$ | m | 0 | -76.4 | -3.3 | 0 | 3625 | -76.4 | 6.56 | -7.48 | $\bar{g}_{HCN,f} = 1e-4$ (soma, dendrite) |
| $I_{HCN,s}$ | $\bar{g}_{HCN,s}m(v - E_h)$ | m | 0 | -87.5 | -4 | 0 | 6300 | -87.5 | 8.9 | -8.2 | $\bar{g}_{HCN,f} = 2.5e-4$ (soma, dendrite) |

**Table 5.** GPe model synaptic current equations.

| Current | Equation | Location | Parameters |
| --- | --- | --- | --- |
| $I_{\text{STN-GPE,AMPA}}$ | $\bar{g}_{syn} s(V - E_{AMPA})$ | distal : $x \geq 100\mu m$ | $\tau_{rise} = 1$<br>$\tau_{decay} = 4$<br>$t_d = 2$<br>$\bar{g}_{syn} = 3.75e-3$ (DD)<br>$\tau_{rec} = 200$<br>$\tau_{facil} = 800$<br>$U_1 = 0.1$ |
| $I_{\text{GPE-GPE,GABA}_A}$ | $\bar{g}_{syn} s(V - E_{GABA_A})$ | proximal : $x < 200\mu m$ | $\tau_{rise} = 2$<br>$\tau_{decay} = 5$<br>$t_d = 0.5$<br>$\bar{g}_{syn} = 2e-4$ (DD)<br>$\tau_{rec} = 400$<br>$\tau_{facil} = 1$<br>$U_1 = 0.2$ |
| $I_{\text{GPE-GPE,GABA}_B}$ | $\bar{g}_{syn} \frac{G^n}{G^n + K_d^n} (V - E_{GABA_B})$ | proximal : $x < 200\mu m$ | $\tau_{rise} = 5$<br>$\tau_{decay} = 25$<br>$t_d = 0.5$<br>$\bar{g}_{syn} = 3e-4$ (DD)<br>$K_3 = 0.098$<br>$K_4 = 6.25e-3$<br>$K_d = 1.4$<br>$n = 4$ |
| $I_{\text{MSN-GPE,GABA}_A}$ | $\bar{g}_{syn} s(V - E_{GABA_A})$ | proximal : $x < 200\mu m$ | $\tau_{rise} =$<br>$\tau_{decay} =$<br>$t_d = 5$<br>$\bar{g}_{syn} = 3e-4$ (DD)<br>$\tau_{rec} = 1$<br>$\tau_{facil} = 200$<br>$U_1 = 0.3$ (DD) |

GPe neurons each had 10 excitatory afferents from STN, 6 inhibitory afferents from GPe-GPe collaterals, and 30 inhibitory afferents from indirect pathway medium spiny neurons (MSN). The glutamergic synapse model was the same as previously described but had only an AMPA component. GABAergic synapses had a fast ionotropic GABA_A_ component with Tsodyks-Markram dynamics, and GPe-GPe synapses had an additional slower metabotropic GABA_B_ component 5.

### Modeling the parkinsonian state

To model the parkinsonian state, the biophysical properties of the network and cell models were modified based on experimental findings reported in the literature. Scaling factors for biophysical parameters were set to experimentally reported values where available, otherwise they were chosen to bring the mean population firing rates into physiological ranges reported for the rat in a state of cortical activation during light anesthesia [2, 53].

The mean firing rate of STN surrogate spike sources was increased from 14.6 to 29.5 Hz in the parkinsonian state [2]. The peak GABA_A_ and GABA_B_ conductance of GPe to STN synapses was increased by 50% and the GABA_B_ decay time constant increased by 2 ms to model the increase in the number of contacts, vesicle release probability, and decay kinetics of GPe afferents [74]. To model the reduction in cortico-STN axon terminals and their dendritic targets [46, 47] the number of CTX-STN afferents was reduced to 70% of the normal condition, corresponding to the ratio of vGluT1 expression in the normal and dopamine depleted condition used to label axon terminals [46]. To model functional strengthening of remaining synapses, the AMPA and NMDA peak conductances of remaining synapses were multiplied by the ratio of the current scaling factors reported in [75] to the fraction of remaining synapses. Finally, HCN currents were reduced by 50% to model reduced depolarization and spontaneous activity after dopamine depletion [76, 77, 77] and modulation of HCN current by D2R receptors [65].

In GPe neurons the peak AMPA conductance of STN afferents was increased by 50% to model the modulatory effect of dopamine on glutamergic excitatory currents [78–80]. The strengthening of GPe-GPe collaterals [81, 82] was modeled by increasing the peak GABA_A_ and GABA_B_ conductances by 50%. The mean firing rate of GPe surrogate spike sources was decreased from 33.7 to 14.6 [9]. Finally, the HCN channel conductance was decreased by 50% in accordance with experimental data [83].

Cortical projection neurons were modeled as Poisson spike generators firing at 10 Hz, a multiple of the experimentally reported rate of 2.5 Hz [84], so that each synapse represented the combined inputs of four pre-synaptic neurons (making use of the additive property of the Poisson distribution). Where cortical oscillatory bursting inputs were modeled, in each period of the beta oscillation 10% of cortical neurons were selected randomly to emit a burst synchronously, using the same phase for the burst onset but a variable number of spikes with inter-spike intervals sampled from the interval [5, 6] ms.

The increase in excitability and spontaneous activity of iMSN [53, 85] was modeled by increasing the mean firing rate of the Poisson spike generators from 1.5 to 6.64 Hz. In experiments where iMSN cells fired oscillatory bursts the same algorithm as described for cortical projection neurons was used. The modulation of GABAergic transmission from iMSN to GPe neurons [86, 87] was modeled by increasing the initial release probability and the peak GABA_A_ conductance of synapses by 50%.

### Simulation details

The model was simulated in the NEURON simulation environment [88] and implemented in Python. The default fixed time step integrator with a time step of 0.025 ms was used for all simulations. Compartmental membrane voltages were initialized to a random value between −63 and −73 mV in GPe and between −60 and −70 mV in STN cells. Gating variables were initialized to their equilibrium values for the initial membrane voltage. Simulation data for the first 2000 ms of each simulation were discarded, and the analyzed intervals were of duration 4000 ms unless otherwise noted. Simulations were run on the UCD Sonic cluster using 8 parallel processes per simulation on a single node, where each node consisted of two Intel Ivybridge E5-2660 v2 CPUs (10 cores per CPU).

### Signal analysis

Signal analyses were performed using the SciPy toolbox [89] for Python. Power spectral densities (PSDs) were calculated using Welch’s periodogram method, using overlapping segments of two seconds duration with 50% overlap and a Hanning window. Given the sampling period of 0.05 ms this led to a frequency resolution of 0.5 Hz. The population PSD was calculated as the mean PSD of all somatic membrane voltages. The instantaneous phase of each population was estimated by applying the Hilbert transform to the average somatic membrane voltage of cells in the population, after band-pass filtering using a neutral-phase filter (Butterworth filter, 4th order, command *sosfiltfilt*) in an 8 Hz wide frequency band centered on the dominant oscillation frequency. For populations that were modeled as surrogate spike trains (cortex and striatum), artificial membrane voltage signals were first constructed by convolving the spike trains with a stereotypical action potential waveform. Bursts were detected using a simple algorithm where a burst consisted of a minimum of four spikes with ISIs ≤ 20 ms.

## Supporting information

**S1 Table.** Experimentally reported connection parameters used to calibrate the model (Table 1).

**S2 Table.** STN neuron model intrinsic current equations from Gillies and Willshaw [43] (Table 2).

**S3 Table.** STN neuron model synaptic current equations (Table 3).

**S4 Table.** GPe neuron model intrinsic current equations from Günay et al. [50] (Table 4).

**S5 Table.** GPe neuron model synaptic current equations (Table 5).

## Acknowledgments

Research supported by the European Research Council (ERC) under the European Union’s Horizon 2020 research and innovation programme (ERC-2014-CoG-646923-DBSModel).

